# The effects of data quantity on performance of temporal response function analyses of natural speech processing

**DOI:** 10.1101/2022.06.07.495139

**Authors:** Juraj Mesik, Magdalena Wojtczak

## Abstract

In recent years, temporal response function (TRF) analyses of non-invasive recordings of neural activity evoked by continuous naturalistic stimuli have become increasingly popular for characterizing response properties within the auditory hierarchy. However, despite this rise in TRF usage, relatively few educational resources for these tools exist. Here we use a dual-talker continuous speech paradigm to demonstrate how a key parameter of experimental design, the quantity of acquired data, influences TRF analyses fit to either individual data (subject-specific analyses), or group data (generic analyses). We show that although model performance monotonically increases with data quantity, the amount of data required to achieve significant prediction accuracies can vary substantially based on whether the fitted model contains densely (e.g., acoustic envelope) or sparsely (e.g., lexical surprisal) spaced features, especially when the goal of the analyses is to capture the aspect of neural responses uniquely explained by specific features. Moreover, we demonstrate that generic models can exhibit high performance on small amounts of test data (2-8 min), if they are trained on a sufficiently large data set. As such, they may be particularly useful for clinical and multi-task study designs with limited recording time. Finally, we show that the regularization procedure used in fitting TRF models can interact with the quantity of data used to fit the models, with larger training quantities resulting in systematically larger TRF amplitudes. Together, demonstrations in this work should aid the learning process of new users of TRF analyses, and in combination with other tools, such as piloting and power analyses, may serve as a detailed reference for choosing acquisition duration in future studies.

## 1. Introduction

Characterizing how acoustic and linguistic features are encoded in human cortex is a key goal of auditory cognitive neurosciences and neurolinguistics. In recent years, substantial progress has been made in utilizing noninvasive electroencephalographic (EEG) and magnetoencephalographic (MEG) responses to continuous speech in order to uncover a diverse set of neural signatures associated with different aspects of language processing. These range from responses to relatively low-level features associated with the speech envelope (e.g., Ding and Simon, 2012; Power et al., 2012; Kong et al., 2014), mid-level features implicated in phonemic processing (e.g., Di Liberto et al., 2015, 2019; Brodbeck et al., 2018), and higher-order linguistic features associated with semantic and syntactic processing (e.g., Broderick et al., 2018; Weissbart et al., 2019; Donhauser and Baillet, 2020; Mesik et al., 2021; Heilbron et al., 2022).

An important catalyst for this work has been the popularization of regularized linear regression methods for mapping the relationship between features in the stimulus space and the brain response (Lalor and Foxe, 2010; Crosse et al., 2016). This relationship can be characterized in both the *forward* direction, mapping from the stimulus to the brain response, and the *backward* direction, specifying how to reconstruct stimulus features from patterns of brain activity. The forward modeling approach models the continuous M/EEG data as a convolution between a set of to-be-estimated feature-specific impulse responses, known as the *temporal response functions* (TRFs), with the known time courses of the corresponding features (e.g., acoustic envelope, lexical surprisal, etc.). These analyses contrast with the more traditional event-related potential (ERP) method, which relies on averaging of hundreds of identical trials in order to estimate the stereotypical neural response for a given stimulus (e.g., Luck, 2005; Woodman, 2010). Unlike ERP methods, which rely on repetition, the TRF approach allows for analyzing brain responses to naturalistic time-varying stimuli, including continuous speech and music.

While the popularity of TRF methods has increased substantially, there remains a relative lack of literature exploring how these methods perform in the context of EEG and MEG analyses under various constraints, such as the type of features utilized (e.g., temporally sparse vs. dense) or the quantity of the data to which the models are applied. This information resource gap has more recently received some attention (Sassenhagen, 2019; Crosse et al., 2021), but published work has provided a broader overview of issues in TRF research (e.g., effects of correlated variables, missing features, preprocessing choices, etc.) without a more thorough examination of any one issue. As such, there is a continued need for further methodological resources aiding researchers interested in adopting TRF techniques to learn using these methods, and to optimize their experimental design for their efficient use.

One of the most fundamental decisions in study design is the choice of how much data to collect (i.e., number of subjects and acquisition duration per subject). This decision has broad consequences affecting the cost of the study, complexity of applicable models, data quality (due to fatigue / comfort level changes over the course of long experimental sessions), and others. With respect to TRF modeling, understanding how analysis outcomes are influenced by data quantity is important for making decisions about duration of data acquisition. Specifically, at the low end of the spectrum (small amounts of data), TRF models may be unable to isolate neural responses of interest due to poor data signal-to-noise ratio (SNR) and/or limited sampling of the stimulus feature space used in the analysis. At the upper end of the spectrum (large amounts of data), model performance may saturate, making additional data wasteful both in research costs and subject discomfort. Understanding these tradeoffs is particularly important for studies of special populations, such as the elderly or clinical patients, who may not tolerate longer experimental sessions.

To date, majority of work exploring effects of data quantity on analyses of speech-evoked EEG data have focused on attention decoding, especially with *backward* models (e.g., Mirkovic et al., 2015; O’Sullivan et al., 2015; Fuglsang et al., 2017; Wong et al., 2018). However, because the driving force behind the interest in attention decoding is innovation in hearing aid technologies, much of this work has focused on the performance of trained models on decoding attention using limited amount of data. In other words, majority of this work has explored the effects of data quantity at the level of model *evaluation,* rather than on the model *training* itself (but see Mirkovic et al., 2015). In the context of forward modeling, the effect of *training* data quantity on model performance has only received a limited amount of attention. Di Liberto and Lalor (2017) explored the impact of data acquisition duration on the performance of TRF models of phonemic processing, to assess whether small amounts of data can reliably support detection of phonemic responses in individuals. They showed that although models trained on data from individual participants required 30+ min to detect these signals, *generic* models derived from data from multiple participants could detect signals related to phonemic processing with as little as 10 min of data per participant. More recently, in their overview of TRF methods, Crosse et al. (2021) used simulations to demonstrate the impact of noise and data quantity on a single-feature TRF prediction accuracies and the fidelity of the derived TRFs. However, beyond these works, a more thorough exploration of TRF forward model performance in the context of more realistic listening scenarios, with a more diverse set of modelled speech features has not been performed thus far.

The goal of the present work is to provide a detailed, practical demonstration of how data quantity and model feature sparsity affect the outcome of TRF analyses in the context of real noninvasive EEG responses to a dual-talker continuous speech paradigm (Mesik et al., 2021). In a series of analyses, we repeatedly fit TRF models to progressively larger segments of the data, estimating both individual subject models, as well as generic models based on data pooled across multiple subjects. For each analysis, we demonstrate how data quantity influences TRF estimates, overall model prediction accuracy, and prediction accuracy attributable to individual features. Additionally, we show the effect of the interplay between data quantity and regularization on the amplitudes of resulting TRF estimates. Given the unique nature of each auditory study (design, analyses, etc.) and the rapid innovation in the TRF methods, we caution readers against taking our work as a sole prescription for how much data should be collected in future TRF studies, or how the resulting data should be analyzed. Instead, we believe our work should serve as a detailed demonstration of general patterns of TRF model performance as a function of data quantity, and a reference that should be carefully used in conjunction with other tools and sources of information (e.g., piloting and power analyses) for informing study design and the subsequent analyses.

## 2. Materials and Methods

EEG data used in the present manuscript was acquired and previously analyzed to investigate age effects in cortical tracking of word-level features in competing speech (Mesik et al., 2021). Extensive description of the details associated with the experiment and the data are openly accessible in the original manuscript. For brevity, we highlight key aspects of this data below.

### 2.1 Participants

Data from 41 adult participants (18-70 years old, mean ± SD age: 41.7±14.3 years; 15 male, 26 female) was used in the present study. The broad age range was utilized to assess age effects on speech-driven EEG responses in the original study. Consequently, a subset of participants (n = 18) had mild-to-moderate hearing loss (HL), largely concentrated in the high-frequency region (≥ 4 kHz), which was compensated for via amplification. All participants provided a written informed consent and received either course credit or monetary compensation for their participation. The procedures were approved by the Institutional Review Board of the University of Minnesota.

### 2.2 Stimuli

Stimuli were four public-domain short-story audiobooks (*Summer Snow Storm* by Adam Chase; *Mr. Tilly’s Seance* by Edward F. Benson; *A Pail of Air* by Fritz Leiber; *Home Is Where You Left It* by Adam Chase; source: LibriVox.org) read by two male speakers (2 stories per speaker). Each story had a duration of approximately 25 minutes. Stories were pre-processed to truncate silent gaps exceeding 500 ms to 500 ms, and the levels in each one-minute segment were root-mean-square normalized and scaled to 65 dB SPL. In participants with HL, the audio was then amplified to improve audibility at frequencies affected by HL (see Mesik et al., 2021 for details of amplification). Stimuli were presented using ER1 Insert Earphones (Etymotic Research, Elk Grove Village, IL).

### 2.3 Experimental procedure

Participants completed two experimental runs in which they listened to pairs of simultaneously presented audiobooks narrated by different male talkers. The stories were presented at equal levels (i.e., 0 dB SNR) and were spatially co-located (i.e., diotic presentation of same stimuli to the two ears). One story was designated as the target story and participants were instructed to attend to the target talker for the duration of the experimental run, while ignoring the other talker. Runs were divided into 1-minute blocks, each of which was followed by a series of four multiple-choice comprehension questions about the target story, along with several questions about the subjects’ state of attentiveness and story intelligibility. This behavioral data was not analyzed in the context of the present manuscript. In the second experimental run, participants listened to a new pair of stories spoken by the same two talkers, with the to-be-attended and to-be-ignored talker designations switched, to eliminate talker-specific effects in the analysis results. The order of the story pairs as well as the to-be-attended talker designations were counter-balanced across participants. All experimental procedures were implemented via the Psychophysics Toolbox (Brainard, 1997; Pelli, 1997; Kleiner et al., 2007) in MATLAB (Mathworks, Natick, MA, United States; version R2019a).

### 2.4 EEG procedure

Data were acquired using a non-invasive 64-electrode BioSemi ActiveTwo system (BioSemi B.V., Amsterdam, The Netherlands), sampled at 4096 Hz. Electrodes were placed according to the international 10-20 system. Additional external electrodes were used to obtain activity at mastoid sites, as well as a vertical electro-oculogram. Data from these external electrodes were not analyzed in the present study.

### 2.5 EEG preprocessing

Here we include a brief overview of pre-processing steps applied to the data. For more detailed description of pre-processing, see Mesik et al. (2021). Unless otherwise stated, pre-processing steps were implemented using the EEGLAB toolbox (Delorme and Makeig, 2004; version 14.1.2b) for MATLAB. To reduce computational load, raw data were downsampled to 256 Hz and band-pass filtered using *pop_eegfiltnew* function between 1 and 80 Hz using a zero-phase Hamming windowed sinc FIR filter (846^th^ order, 1 Hz transition band width). Next, the data were pre-processed using the PREP pipeline (Bigdely-Shamlo et al., 2015), in order to reduce the impact of noisy channels on the referencing process. This procedure involved three stages: 1) power line noise removal via multi-taper regression, 2) iterative referencing procedure to detect noisy channels based on abnormally high signal amplitude, abnormally low correlations with neighboring channels, poor predictability of channel data based on surrounding channels, and excessive amount of high-frequency noise, and 3) spherical-interpolation of the noisy channels detected in stage 2. For this procedure, we used the default parameters outlined in Bigdely-Shamlo et al. (2015). Following up to 4 iterations of stages 2-3 (or until no further noisy channels were identified), the cleaned estimate of global mean activation was used to reference the dataset.

Subsequently, we epoched all 1-minute blocks and applied independent component analysis (ICA; Jutten and Herault, 1991; Comon, 1994) to remove components of data corresponding to muscle artifacts and other sources of noise. ICA decomposes EEG signal into a set of statistically independent components that reflect various underlying contributors to the channel data (e.g., eye blinks, different aspects of cognitive processing, etc.), allowing for removal of components driven largely by nuisance factors such as muscle activity.

The data were then band-pass filtered between 1 and 8 Hz using a Chebyshev type 2 filter (80 dB attenuation below 0.5 Hz and above 9 Hz, with 1 dB band-pass ripple), applied with the *filtfilt* function in MATLAB. This was done given the existing evidence that cortical speech processing mechanisms track speech predominantly via low-frequency dynamics in the 1-8 Hz range (e.g., Ding and Simon, 2012; Zion Golumbic et al., 2013). Finally, the data were transformed into z-scores to account for variability in overall response amplitudes due to inter-subject variability in nuisance factors such as skull thickness. Data from the first block of each run were excluded from analysis due to a small subset of participants accidentally confusing the attended and ignored speakers in the initial block (this became apparent during behavioral task following the first block, which pertained to the to-be-attended story).

### 2.6 TRF analyses

The time courses of speech-evoked responses, or TRFs, were extracted from the pre-processed EEG data using regularized linear regression (i.e., ridge regression), implemented via the mTRF Toolbox (Crosse et al., 2016, version 2.3). Briefly, TRFs are estimated by regressing a set of *n* time-lagged copies of the time series of a given speech feature (e.g., acoustic envelope) against the EEG time course at *m* channels. This results in *m* separate TRFs, each representing how the response at a given electrode site is affected at the *n* time lags relative to that feature’s presentation times. This procedure can be simultaneously applied to multiple features (see section 2.6.3 for details of features used in our analyses), enabling the decomposition of EEG signals into contributions from different stages of speech processing (e.g., acoustic vs. semantic processing). Due to the non-white spectral content of speech signals, the TRF estimation procedure is prone to high degree of variance in the resulting TRFs (Crosse et al., 2016). Regularized (ridge) regression introduces a “smoothing” bias into TRF estimation by penalizing large regression coefficients, resulting in the reduction of this (undesirable) variance. Note that other methods of TRF estimation procedures, such as “boosting” (David et al., 2007; Brodbeck et al., 2021), rely on alternative approaches to regularization. These alternative methods were beyond the scope of this work, although recent work directly comparing these methods suggests that they typically produce highly similar results (Kulasingham and Simon, 2022).

To minimize overfitting, the regression procedure was implemented using leave-one-trial-out cross-validation in which all-but-one (training) trials or subjects (see sections 2.6.1 and 2.6.2 for details) were used to estimate the TRFs, and the remaining held-out (test) trial/subject was used to evaluate the prediction accuracy of the estimated model parameters. In the fitting stage of each cross-validation loop, the data were first used to select the optimal regularization (ridge) parameter λ. This was done via a separate leave-one-out cross-validation loop utilizing only the training trials. In each of these cross-validation folds, the model was fit using a range of different λ parameter values and evaluated by predicting the data from the left-out trial. The prediction accuracies for each λ value were then averaged across all cross-validation folds and electrodes. The λ corresponding to the highest average prediction accuracy was used in the final fit to the entire training set. The resulting fit was then evaluated on the held-out “test” trial or subject to determine the ability of the TRF model to predict EEG responses to speech. The Pearson’s correlation between the predicted and the actual data represents the prediction accuracy. This procedure was repeated *j*-times, where *j* denotes the number of trials or subjects, each time holding out data from a different trial/subject, resulting in *j* sets of TRF estimates and prediction accuracies.

In addition to overall model prediction accuracies, we further estimated the degree to which each feature contributed to the model performance. Briefly, this was done by comparing the full model’s prediction accuracy to “reduced” model prediction accuracies, which were obtained by fitting models in which individual features were excluded. For example, to estimate contribution of feature “A” in a model containing features “A” and “B”, we computed the prediction accuracy difference between model containing both features, to model containing only the feature “B”. In other words, the additional predictive power of a full model over a reduced model can be interpreted as reflecting the contribution of the extra feature present in the full model. In the remainder of this manuscript, these differentials are referred to as feature-specific model contributions.

The main goal of the present work was to explore how estimates of cortical responses to a range of speech features are influenced by the quantity of EEG data included in the analysis. To explore this question, we iteratively applied identical sets of analyses to progressively larger amounts of the pre-processed EEG data. This was done via two distinct analysis approaches: 1) subject-specific analyses, where data from each participant was fit independently, and 2) generic analyses, where data from multiple subjects were fit jointly. Below we provide details of each analysis approach.

#### 2.6.1 Subject-specific analyses

The subject-specific analyses involved repeated TRF estimation using data from individual subjects with progressively larger amounts of their data. Cross-validated fitting procedure was repeated with 11 distinct data quantities (3, 4, 6, 8, 10, 14, 18, 24, 30, 36, and 42 min of data), with data in each analysis selected in chronological order to reflect real constraints of data collection. While this may potentially bias analysis outcomes via temporally systematic phenomena such as fatigue or adaptation, such order effects are a natural aspect of most experiments and thus reflect realistic data acquisition scenarios. The data quantities in this, as well as generic analyses (see section 2.6.2), were chosen to be spaced quasi-logarithmically to more densely sample changes in model performance at relatively low data quantities, under the assumption that at larger data quantities performance would begin to saturate. Group-level analyses were performed using each subject’s average TRFs and prediction accuracies across all cross-validation folds (see section 2.6.4).

#### 2.6.2 Generic subject analyses

In generic subject analyses, we repeatedly estimated TRFs using data pooled across progressively larger number of subjects. Cross-validated model fitting was again repeated with 11 distinct numbers of subjects (3, 4, 6, 8, 10, 14, 18, 24, 30, 36, and 41 subjects) at three distinct data acquisition durations per subject (2, 4, and 8 min), yielding 33 unique analysis outputs per model. The data per subject was constrained both because of memory limitations for analyses involving larger numbers of subjects, but also to explore the extent to which small amount of data per subject can support accurate TRF estimation and robust prediction accuracies.

A notable limitation of generic analyses, as implemented here, is that all of the data is utilized within a single cross-validation procedure, resulting in a single set of average prediction accuracy and TRF estimates. While individual subject prediction accuracies from cross-validation can be used for group level statistics, these statistics are highly susceptible to noise from outlier data when utilizing small numbers of subjects. As such, to assess the central tendencies of generic analysis performance more accurately as a function of subject count, we utilized a resampling approach to obtain distributions of model performances for each sample size. Briefly, for each analysis (i.e., sample size), we randomly resampled, with replacement, participant data 20 times, and used leave-one-subject-out cross-validation procedure to fit each such resampled data set with a TRF model. We then used the mean TRFs and prediction accuracies across the cross-validation folds for further analyses, yielding 20 sets of TRFs and prediction accuracies for each sample size, model, and per-subject data quantity. Note that the 20 resamplings were performed independently at each sample size, rather than being done in a dependent manner via incremental sampling (with increasing sample size) of more participants into the to-be-analyzed data pool. To obtain maximally comparable full and reduced model performances for estimating feature-specific model contributions (see section 2.6, paragraph 3), we utilized identical sets of resampling indices (i.e., same participant data) for full and reduced models.

To aid statistical evaluation of the analyses (see section 2.7.2) we corrected the model prediction accuracies using estimates of the noise floor, i.e. range of prediction accuracies that may be expected by chance, by mismatching regressor-data pairings and computing their corresponding prediction accuracies. This was done at the level of each cross-validation fold (i.e., held-out subject), where we computed prediction accuracies for each 1-min data segment for all mismatched regressor-data pairings. Each subject’s true prediction accuracy was then noise-floor corrected by subtracting the average mismatched prediction accuracy. Results of the 20 resampling analyses were then interpreted as a distribution of mean TRFs and prediction accuracies expected for a given sample size. Because each of the 20 iterations corresponded to 5% of the analyses, results in which all 20 analyses showed consistent results (e.g., same sign of prediction accuracies) were interpreted as statistically significant (albeit at a level uncorrected for multiple comparisons). Note that to allow for more reliable statistical inference, bootstrap-based analyses such as these are typically performed ≥1000 times in order to more precisely estimate the degree of overlap between the distributions of parameter estimates for different conditions. However, because the goal of this work was to demonstrate general behavior of TRF analyses rather than to draw strong statistical conclusions about our data, and the computational load of running ≥1000 resamplings for each of the 66 analyses (2 models x 11 subject counts x 3 data quantities/subject) would have been very high, we chose to perform the more modest bootstrap procedure with 20 iterations.

#### 2.6.3 Modelled speech features

To explore effects of feature choice on subject-specific and generic model performance, each modeling approach was evaluated using two distinct models of *attended* speech processing that emphasized features with different levels of sparseness. In the denser “envelope” model we included a continuously varying log-transformed acoustic envelope, along with a sparse word-onset regressor intended to capture responses to acoustic onsets and lexical segmentation. The sparser “surprisal and SNR_word_” model included lexical surprisal for each word, SNR of each attended word against the to-be-ignored speaker, and the word-onset regressors. While lexical surprisal has been shown to be robustly tracked by cortical language processing mechanisms (e.g., Weissbart et al., 2019; Heilbron et al., 2022), SNR_word_ is a relatively unexplored feature in the context of TRF literature. We chose to include it to reflect the fact that the attended talker’s intelligibility fluctuated on a moment-to-moment basis due to the independent variations in the relative attended and ignored talkers’ speech intensities. The “glimpsing” models of speech understanding have proposed that in speech-on-speech masking scenarios, time periods with relatively high SNR (i.e. “glimpses”) may serve a privileged role in speech perception, with information from glimpsed periods being used to fill-in information from masked portions of speech (e.g., Cooke, 2006).

Word-onset regressors were included in both models primarily to help account for onset-driven neural activity, which is known to have particularly large amplitudes. While speech signals generally contain numerous onsets beyond those found at word onsets, these occur largely at faster rates (up to 50 Hz; e.g., Stone et al., 2010) than the 1-8 Hz range analyzed here. As such, we abstained from utilizing envelope onset regressors (e.g., half-wave rectified envelope derivative), as they would fall outside the passband of low-rate envelope representations, and thus would not contribute to model prediction accuracies. This was confirmed in a supplemental analysis, where envelope onset representations did not provide additional predictive power in the 1-8 Hz frequency band (results not shown).

Besides their level of sparseness, the two feature sets also differed in terms of the modelled response latencies, as envelope responses have previously been shown to occur largely at relatively low latencies < ~300 ms (e.g., Power et al., 2012; Kong et al., 2014; Fiedler et al., 2019), while responses to sparser, higher-order features occur at longer latencies (e.g., Weissbart et al., 2019; Mesik et al., 2021). Responses to features in the denser model were therefore modelled using time intervals between −100 to 350 ms relative to feature onsets, while responses to the features in the sparser model were modelled between −100 to 800 ms relative to feature onset. Consequently, fitting the denser model to signal recorded by a particular electrode resulted in estimation of 234 model beta coefficients (2 features × 117 samples, corresponding to the time lags above), whereas the sparser model resulted in 696 beta coefficients (3 features × 232 samples).

Across the two models, the four features were estimated as follows. The word onset regressors contained unit-amplitude impulses time-aligned to the onset of each word. The low frequency acoustic envelopes were extracted by half-wave rectifying the speech stimuli and lowpass filtering this representation below 8 Hz. This representation was then log-transformed to more closely approximate encoding of sound level in the human auditory system. Lexical surprisal regressors were the negative logarithm of each word’s probability given the multi-sentence preceding context, as estimated using the GPT-2 artificial neural network (Radford et al., 2019). SNR_word_ regressors were obtained by computing the ratio of each word’s root-mean-square (RMS) of its acoustic waveform and the RMS of background speaker’s acoustic waveform. This was motivated by the increased importance of momentary glimpses of high broadband SNR for speech understanding at low target-to-masker ratios (Best et al., 2019). As with envelopes, these ratios were log-transformed to more closely mimic sound level encoding in the human auditory system. Features in both surprisal and SNR_word_ regressors utilized the same timing as word-onset regressors, as prior work has shown that onset timing results in reasonable characterization of responses to higher-order speech features (e.g., Broderick et al., 2018; Weissbart et al., 2019; Mesik et al., 2021). Finally, non-zero regressor values for all features were RMS scaled to have an RMS value of 1. This was done to make TRFs for different features more similar in amplitudes in order to optimize regularization performance. With the exception of the acoustic envelope, a more detailed description of the derivation of these features can be found in Mesik et al. (2021).

Note that different features were separated into the denser and sparser models for the practical purpose of exploring how performances of models with distinct levels of sparseness vary as a function of data quantity used in fitting. However, outside of this context, pooling features into larger models (e.g., containing both acoustic and linguistic features) can generally be advantageous, especially for more accurate estimation of feature-specific model contributions (e.g., Gillis et al., 2021; see section 4.4.5).

#### 2.6.4 Regions of interest

Due to the relatively low spatial resolution of EEG data and for simplicity of analysis result presentation, we chose to limit the spatial dimensionality of the results to two regions of interest (ROIs), the frontal and parietal ROIs. These ROIs were chosen based on the peak locations of activation for the two models. Both ROIs contained 13 electrodes, which symmetrically surrounded electrode Fz in the frontal ROI (i.e., AF3, AFz, AF4, F3, F1, Fz, F2, F4, FC3, FC1, FCz, FC2, and FC4), and electrode Pz in the parietal ROI (i.e., CP3, CP1, CPz, CP2, CP4, P3, P1, Pz, P2, P4, PO3, Poz, and PO4). ROI-specific TRFs and prediction accuracies were computed by averaging TRF analysis results from these sets of electrodes. All statistical analyses were performed on these ROI-averaged results. Note that the choice of ROI-based analyses was driven primarily by the simplicity of data presentation, for educational purposes. TRFs of individual features may potentially perform better at specific electrode sites not directly analyzed here.

### 2.7 Statistical analysis

#### 2.7.1 Statistical analyses of Subject-specific models

The primary goal of this work was to describe general patterns of TRF model behavior as a function of data quantity used in model fitting. For subject-specific analyses, to test whether a given quantity of data was sufficient for the derived TRF to yield prediction accuracies that were significantly greater than zero, we utilized either t-tests or Wilcoxon signed-rank test, based on the outcome of the Anderson-Darling test of normality. These tests were conducted at the group level, using all 41 individual prediction accuracies. To normalize the distribution of correlation coefficients, their values were Fisher z-transformed prior to conducting the statistical tests. Note that statistics were not corrected for multiple comparisons, as we treated the analysis of each data quantity as a quasi-independent experiment, emulating the scenario where only that amount of data was acquired. Because analyses on different data quantities were not independent due to utilizing partially overlapping data, we abstained from direct pairwise comparisons of model prediction accuracies, and instead focused on more general description of model performance patterns (e.g., trajectory of mean model performance and between-subject variance) as a function of data quantity. However, in order to provide a statistical description of linear changes in model performance as a function of data quantity, we fit linear mixed effects models (LMM) to the prediction accuracy results (as well as each feature’s unique model fit contributions), with a fixed effects of training data quantity, ROI, and a random effect of participant identification number. In all LMM analyses, training data quantity was treated as a continuous numeric variable, while ROI was treated as a categorical variable. Significance of these models were assessed using LMM ANOVA (type 3) and Satterthwaite approximation for degrees of freedom.

We additionally used the subject-specific model fits to explore the relationship between the size of participant pool and the data quantity per subject required to achieve statistically reliable detection of cortical tracking of attended speech for different significance levels. To do this, we utilized subject pool sizes ranging from 2 to 41 subjects, and for each pool size, we resampled with replacement the prediction accuracies from analyses of each training data quantity (i.e., minutes of data per participant) 10,000 times. Within these samples, we conducted t-tests on results derived from each of the 11 data quantities per subject, searching for the minimum data quantity for which at least 80% of the 10,000 analyses exceeded the t-score thresholds corresponding to p < 0.05, p < 0.01, and p < 0.001. For these analyses we used parametric statistics, although non-parametric tests resulted in similar patterns of results.

#### 2.7.2 Statistical analyses of generic models

For generic analyses, we estimated prediction accuracy noise floors as described in section 2.6.2. Within each cross-validation fold of each resampling analysis, we computed the difference between the true prediction accuracy and the “mismatched” prediction accuracy estimated using mismatched regressor-data pairings for the held-out participant. Thus, the distribution of these corrected prediction accuracies across resampling analyses reflects the proportion of times in which the true regressor-data pairings enabled more accurate data predictions than mismatched-pairings. Given that our analyses included 20 resamplings, only analyses where all data points exceeded the 0-point were deemed to be significant (i.e., since 1/20 corresponds to 5%). Note that noise floor correction in generic analyses was motivated by the possibility that very small positive prediction accuracies could occur by chance, making it difficult to determine the proportion of analyses with reliably elevated prediction accuracies. This correction was particularly important given our use of only 20 resampling analyses. In principle, this approach could also be used in subject-specific analyses. However, we abstained from the use of noise floors in subject-specific analyses because of their greater statistical power (i.e., more independent data points), our use of cross-validation, and the observation that in generic analyses, noise floor prediction accuracies were concentrated around zero.

To test for linear increase in prediction accuracy as a function of number of participants used in generic analyses, we fit linear functions to the patterns of generic model prediction accuracies. Due to the dependent nature of the 20 resampled estimates of prediction accuracies at each training data quantity, we used a separate bootstrap analysis approach to establish the likelihood of observing a positive slope relating training data quantity and prediction accuracies. Specifically, on each bootstrap iteration we randomly selected one result from the pool of resampling analyses for each training data quantity, and fit a two-parameter linear function (slope and intercept) to the obtained series of 11 prediction accuracies. We repeated this procedure 1000 times, each time storing the slope of the linear fit. Finally, we deemed the result of each such analysis significant if >95% of obtained slopes were greater than zero. This procedure was done separately for noise floor-corrected overall prediction accuracies at each training data quantity per subject, as well as the corresponding feature-specific model contributions, of both the denser and sparser models.

For each model type and feature set, we assessed the similarity of TRFs derived using different data quantities by computing pairwise Pearson’s correlations between time courses of the TRFs. Additionally, we assessed the extent to which subject-specific and generic analyses resulted in estimation of morphologically similar TRFs by computing Pearson’s correlations between these TRFs. For these analyses, we focused on models that utilized the greatest amounts of minutes (subject-specific analyses) and subjects (generic analyses). Because edge artifacts are commonly observed in beginning and end TRF samples, all TRFs were trimmed by 25-ms at each end when computing TRF similarities, as well as for visualization purposes.

## 3. Results

We explored the effects of training data quantity on performance of TRF models trained and evaluated on data from individual subjects (section 3.1), or trained on aggregate data of multiple subjects and evaluated on different subjects (section 3.2). Below we discuss each modeling approach in terms of its prediction accuracies, feature-specific model contributions, and the recovered TRFs.

### 3.1 Subject-specific analyses

In subject-specific analyses, each participant’s data was fit individually using each of the two models and evaluated on held-out data from the same participant. Group-level pattern of overall model prediction accuracies as a function of data quantity is depicted in Fig 1 for the denser (Fig 1A) and sparser (Fig 1B) models. In general, increases in training data quantity resulted in monotonic increases in performance for both models (LMM ANOVA: effect of data quantity at p < 0.001 for both models; See Supplementary Table 1), along with reductions in inter-subject variability in prediction accuracies. With 41 participants used in these analyses, both models reached high degree of statistical significance with as little as 5 minutes of data per participant. However, at low training data quantities (e.g., < 10 min), a subset of participants exhibited prediction accuracies (i.e., correlations between the predicted and actual EEG data) that did not exceed 0. At larger data quantities (e.g. > 20 min), prediction accuracy distributions were elevated and largely did not span the value of zero.

**Figure 1.**
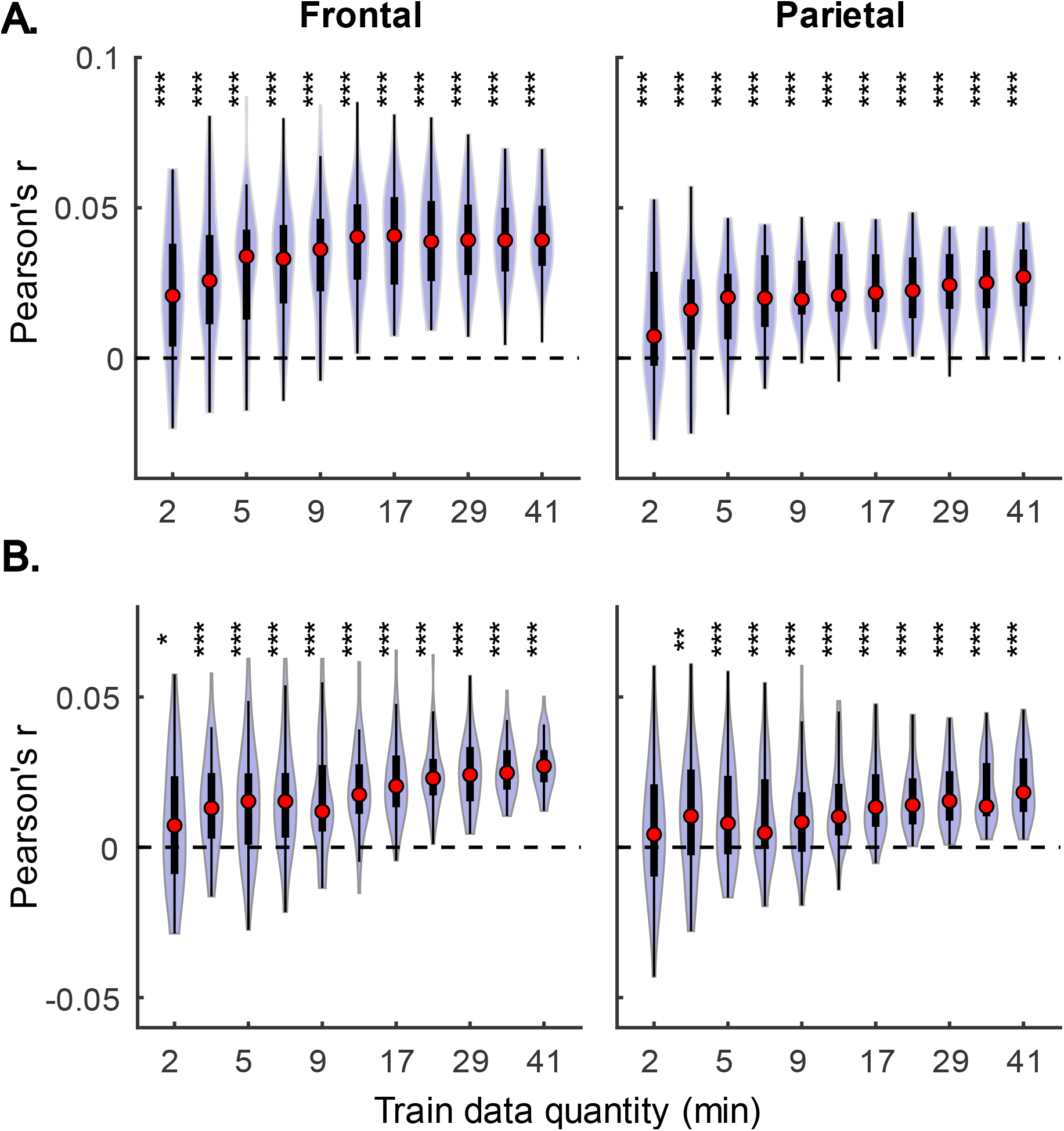
Overall subject-specific model prediction accuracies for the denser (A; onset and envelope features) and sparser (B; onset, surprisal, and SNR_word_ features) models displayed as a function of training data quantity (data acquisition duration). The violin plots depict the distribution of these values for all 41 participants, with a box plot depiction (interquartile range) shown within each violin. Red circles denote median values of the distributions. The uncorrected significance levels of each statical comparison against zero are displayed at the top of each plot: * p < 0.05, ** p < 0.01, *** p < 0.001

Although overall prediction accuracy is a key metric in evaluating TRF model performance, it does not reflect the extent to which individual features in the model uniquely contribute to this accuracy. To assess the unique contributions from individual features, we compared the full model prediction accuracies to those estimated using reduced models containing all-but-one feature. Model fit contributions from individual features (i.e., change in prediction accuracy due to adding a given feature) as a function of training data quantity are depicted in Fig 2 for the denser and Fig 3 for the sparser models, respectively. In contrast to overall prediction accuracies, feature-specific contributions showed a less pronounced increase as a function of data quantity used in model fitting (see Supplementary Table 2), especially in the case of the sparser model, although generally models trained with more data allowed for more robust detection of unique contributions of individual features. Notably, the dense envelope feature (Fig 2, bottom) had substantially higher model prediction accuracies than either the onset feature in the dense model, or any of the sparse features (Fig 3). Accordingly, highly significant detection of unique contribution of acoustic envelope was possible with mere minutes of data. In the sparser model, highly significant model fit contributions were observed only at relatively large training data quantities (> ~17 min) for the word onset feature frontally, and lexical surprisal parietally. For SNR_word_, weakly significant contributions were observed sporadically, with no systematic relation to data quantity. In fact, at very low data quantities, inclusion of SNR_word_ in the model led to a small decrease in prediction accuracies, likely due to the full model having too many parameters relative to the training data quantity. As such, with the available data quantity and the chosen method for estimating feature-specific model contributions, we were only able to detect reliable model contributions for two out of three features of the sparser model.

**Figure 2.**
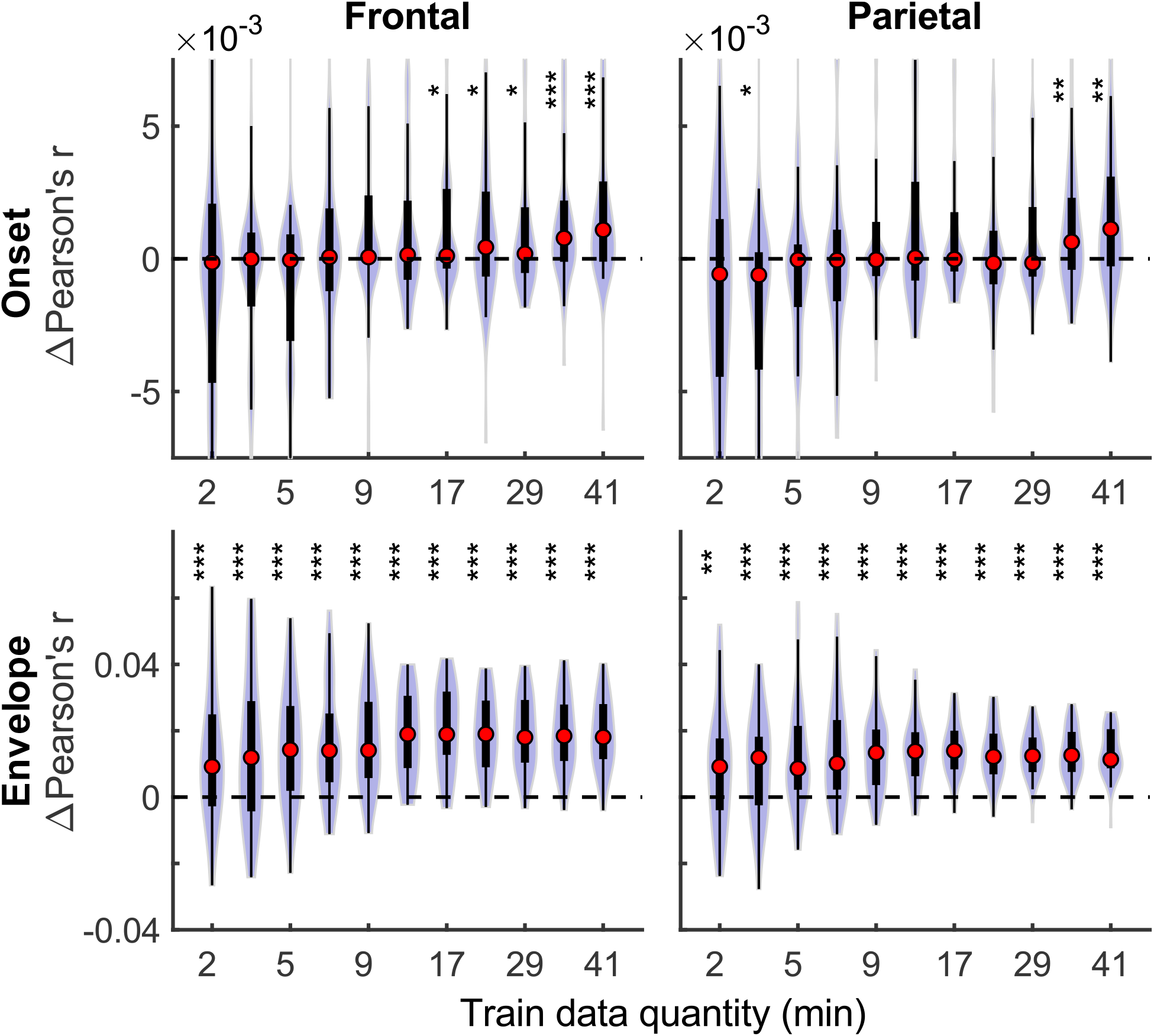
Feature-specific model fit contributions as a function of training data quantity for the denser model. Model fit contributions reflect the unique contribution of each feature to the overall prediction accuracy of the full model, compared to reduced models in which those features were excluded. Note the different scales of the contributions for the Onset (upper row) and Envelope (lower row) features, with the latter fit contributions being about an order of magnitude larger than the former. To facilitate visualization of the central portion of the distributions, upper and lower limits of some of the violin plots for the “Onset” feature are truncated. Asterisks denote significance levels as in Fig 1.

**Figure 3.**
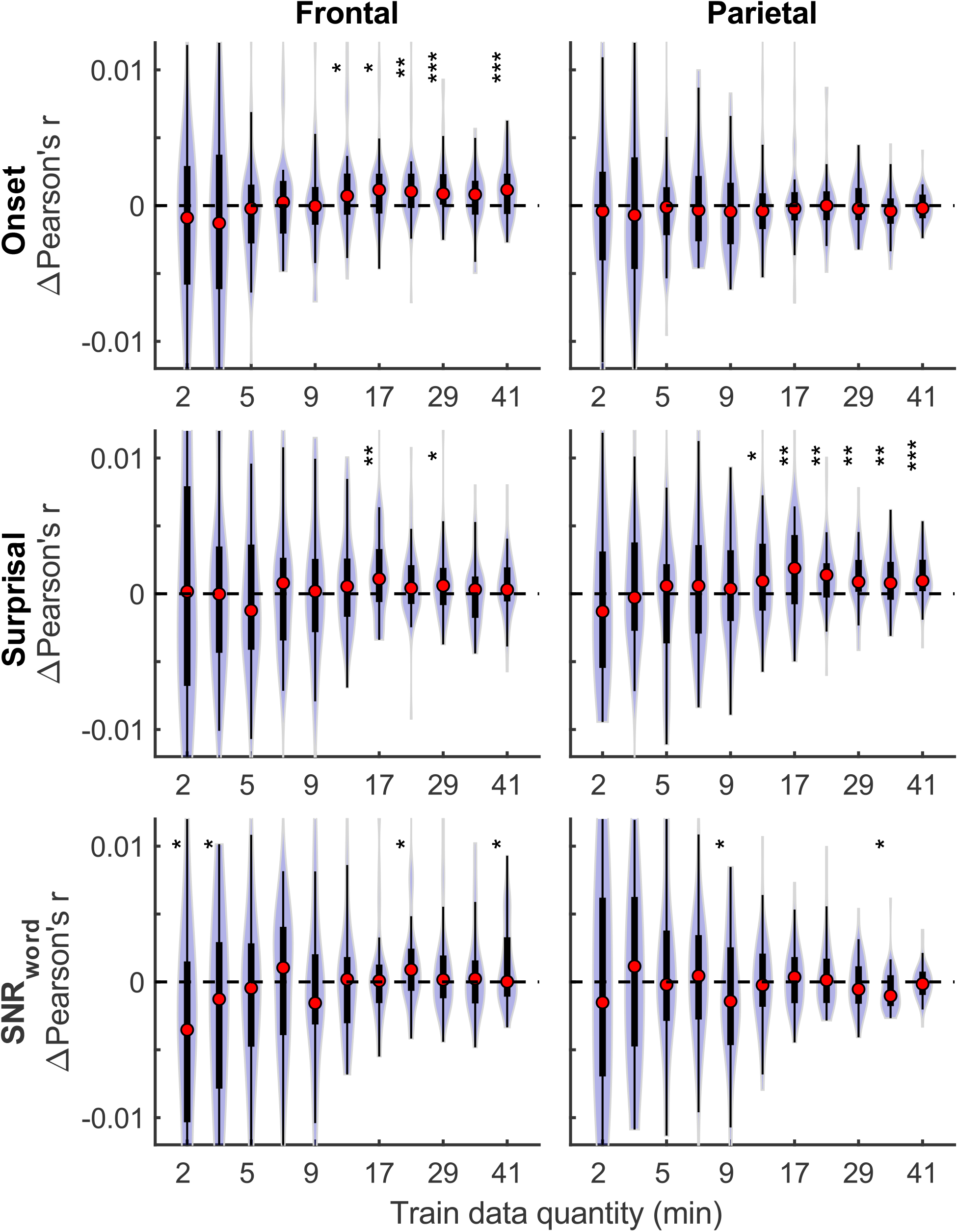
Feature-specific model fit contributions as a function of training data quantity for the sparser model. The data presentation and significance markers are as in Fig 2.

In addition to prediction accuracies, a key output of TRF models are the TRFs themselves: the impulse responses to each of the modelled features. Fig 4 shows TRFs for the denser model and Fig 5 shows TRFs for the sparser model. Mirroring the high prediction accuracies and feature-specific model contributions, the denser model TRFs (Fig 4) showed a high degree of morphological similarity across the different data amounts, albeit with a systematic increase in amplitudes seen with increasing data quantity (for easier comparison across data quantities, see Supplementary Fig S1). In line with its lower feature-specific model contributions, the sparser model TRFs (Fig 4; Supplementary Fig S2) generally exhibited greater noisiness at low data quantities, with TRFs reaching stable form once substantial data quantity (> 17 min) was used in model fitting. These qualitative observations were supported using correlational comparisons of TRF time courses derived using different training data quantities (see Supplementary Figs S3-4 for correlation matrices for denser and sparse models, respectively). While there was a trend for lower TRF amplitudes with increasing data quantity, this pattern is opposite to that seen for the denser model and generally appears less systematic than that seen in Fig 4. Additionally, for both the denser and sparser models, RMS amplitudes of the TRFs showed significant positive correlations with the feature-specific model contributions, particularly in models trained on large data quantities (data not shown). This supports the often assumed association between larger TRF amplitudes reflecting higher predictive power.

**Figure 4.**
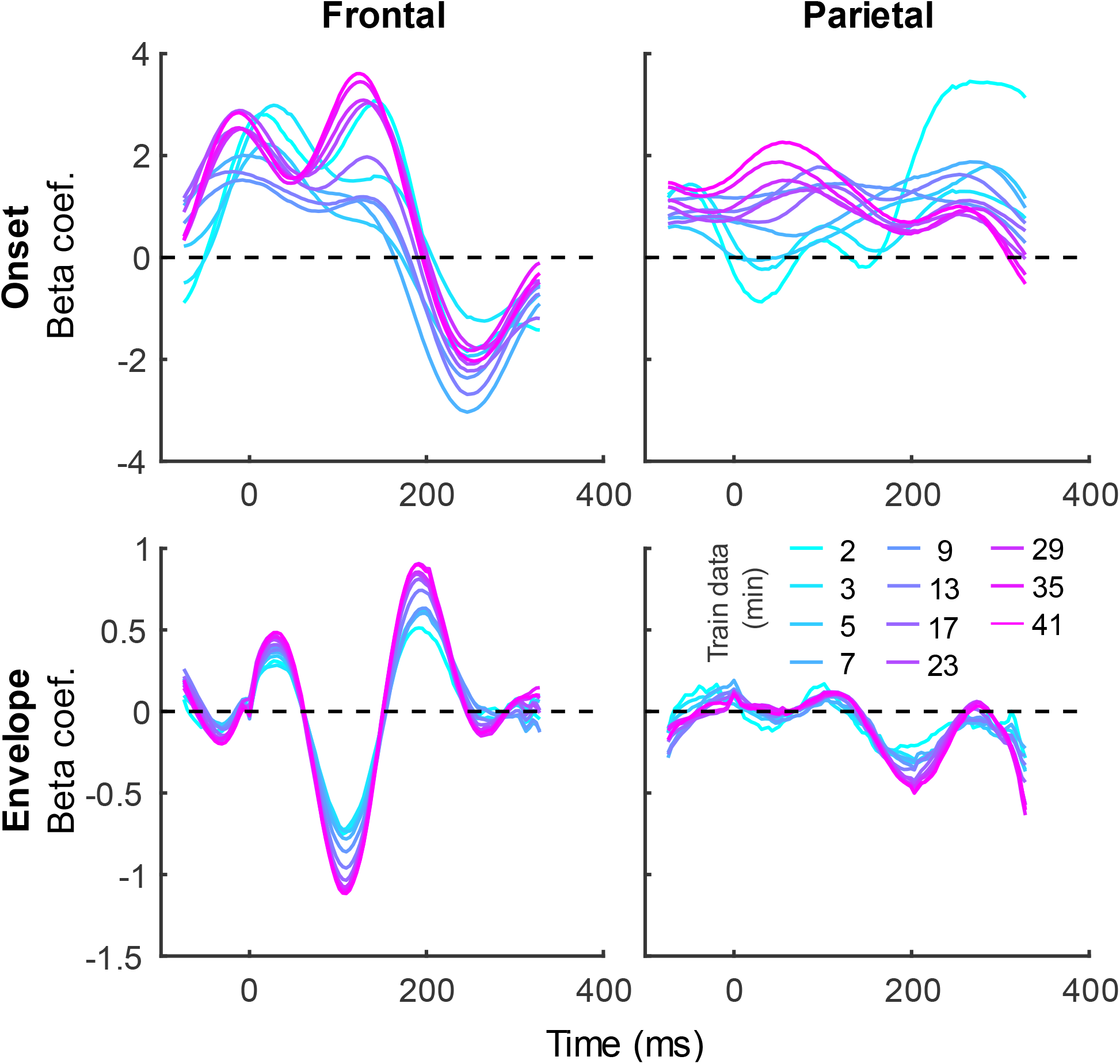
Group-averaged TRF time courses of model beta coefficients for onset (top row) and envelope (bottom row) features from the denser model in frontal (left column) and parietal (right column) ROIs. TRFs estimated using different amounts of data are depicted in different colors (see legend above the lower right plot). The small sharp deflections in envelope response around t = 0 ms reflect mild leakage of electrical artifact from earphones into the EEG signal. See also heatmap depiction of the TRFs in Supplementary Fig S1 for easier comparison across different data quantities, and Fig S3 for correlation matrices comparing TRF morphologies across different data quantities.

**Figure 5.**
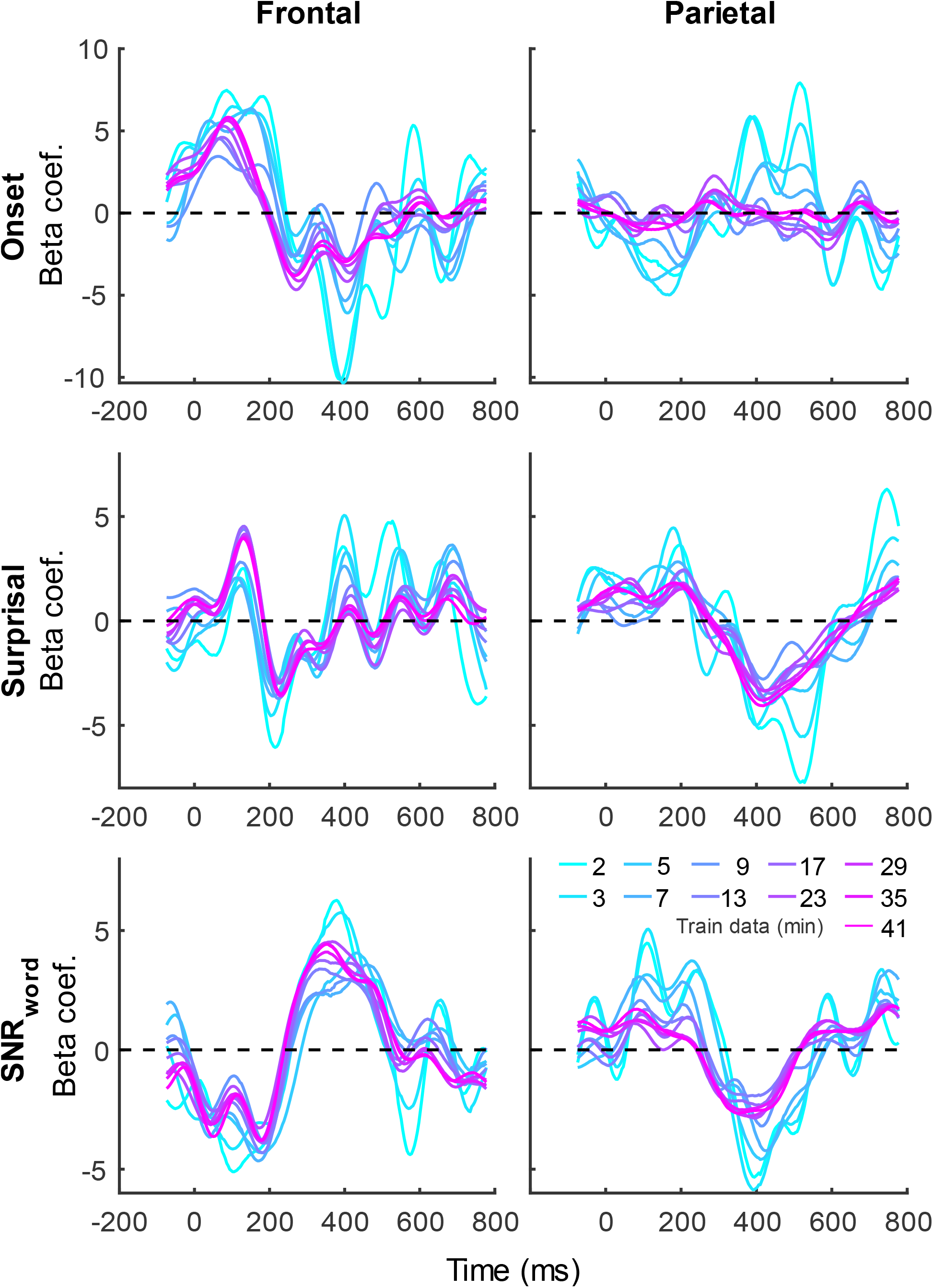
Group-averaged TRF time courses of model beta coefficients for onset (top row), surprisal (middle row), and SNR_word_ (bottom row) features from the sparser model in frontal (left column) and parietal (right column) ROIs, as a function of training data quantity (see legend in bottom right). Note that these TRFs contain a broader range of latencies than those in Fig 3 due to these features engaging higher-order processing, reflected in key TRF features such as the N400 response seen in parietal ROI. See also heatmap depiction of the TRFs in Supplementary Fig S2 for easier comparison across different data quantities, and Fig S4 for correlation matrices comparing TRF morphologies across different data quantities.

The systematic increase in TRF amplitudes seen in Fig 4 could, in principle, reflect a genuine neural phenomenon related to the chronological inclusion of data into models trained on more data. For example, it could reflect improved neural tracking of the target audiobook over the course of the study session. However, examination of the regularization parameter (Fig 6A), which controls the penalty assigned to large TRF values during the fitting procedure, indicates a systematic decrease in this parameter with increasing data quantity. In other words, models fitted to greater data quantity penalized large TRF amplitudes less, likely resulting in the systematic increase in their amplitudes seen in Fig 4. This interpretation was supported by a supplemental analysis (not shown), in which we fit the denser TRF model using a 6-min moving window to assess whether the TRF amplitude changes over the course of the study session. This analysis revealed virtually identical TRF amplitudes in all analysis windows, lending support to the effect in Fig 4 being entirely technical in nature. Regularization parameter of the sparser model (Fig 6B) showed a similar systematic decrease with increasing data quantity without the corresponding increase in TRF amplitudes (Fig 5). This may reflect the overall greater noisiness in these TRFs at low data quantities (see Supplementary Fig S4), as well as greater similarity in general TRF amplitudes across the three features (in contrast to the larger amplitude discrepancy between onset and envelope TRFs seen in Fig 4).

**Figure 6.**
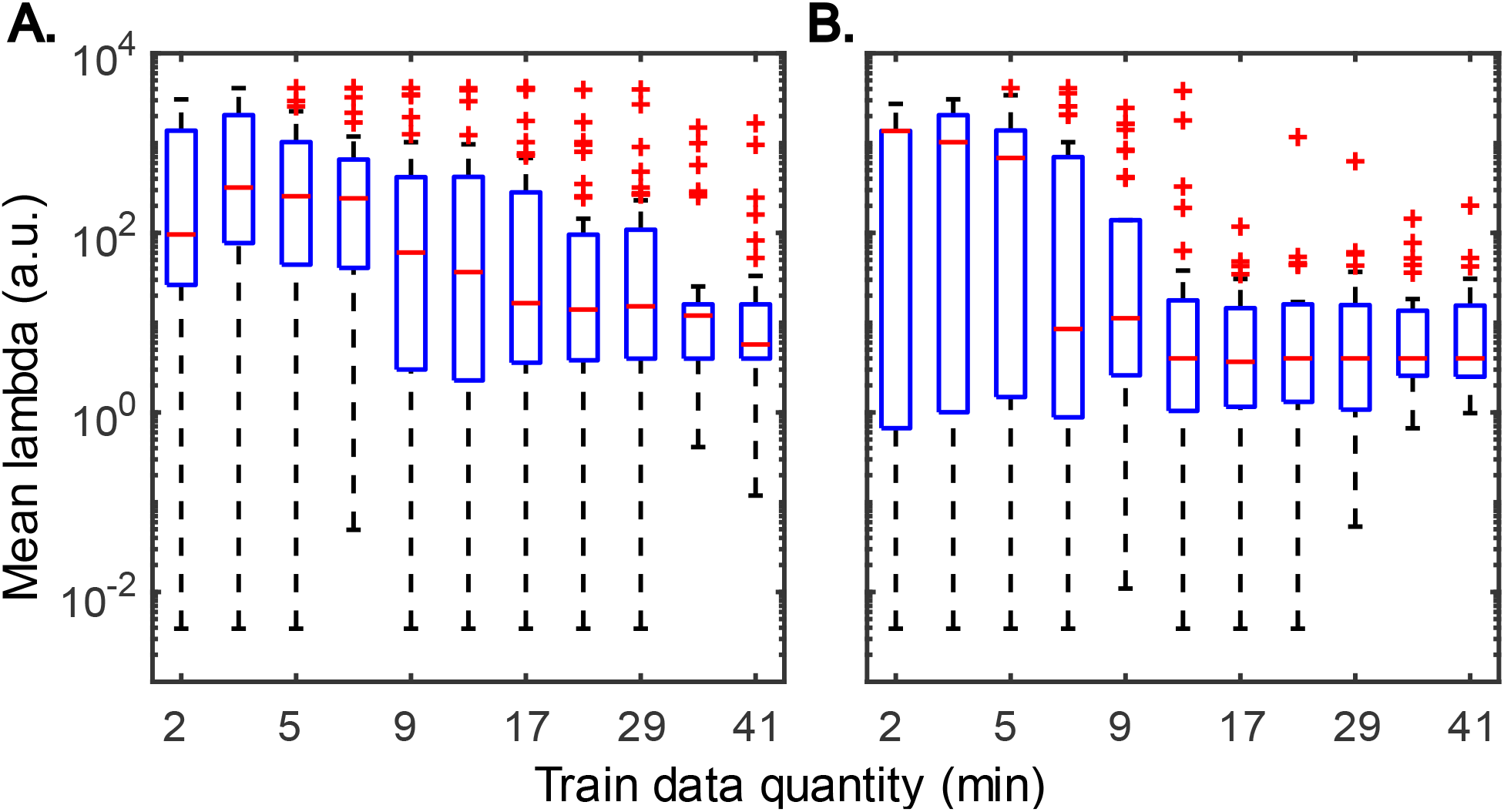
Distributions of subject-specific regularization parameters, lambda, from the denser (A) and sparser (B) models, as a function of training data quantity. Red horizontal lines within each box represent the median value while the lower and upper bounds of the blue boxes depict the 25^th^ and 75^th^ percentiles of the distributions. To aid the visualization of where bulk of the data points were, unusually large parameters (for a given data quantity) were marked as outliers with red “+” symbols. Note: “a.u.” denotes arbitrary units.

Although utilization of data from all participants for analyses in Figs 1–6 provides the most accurate estimates of the average model performance for different training data quantities, it is less informative about the statistical performance of our TRF models with more limited samples. To provide a more complete description of how subject-specific models perform under more limited sample sizes, we performed a resampling analysis using subsets of participant from our 41-subject pool. More specifically, for each participant pool ranging from 2-41 participants, we resampled (with replacement) the subject-specific result pool 10,000-times at each of the data quantities and determined how many minutes of data per participants were required to reach significance in at least 80% of analyses at three commonly used significance thresholds (p < 0.05, 0.01, and 0.001). The results of these analyses are shown in Fig 7 for overall prediction accuracy, and Fig 8 for feature-specific model contributions. These results show the expected downward sloping pattern whereby larger participant pools require smaller amount of data per participant. Additionally, these results mirror those in Figs 1–3 in that the sparser model generally requires more data per participant, and capturing significant feature-specific model contributions requires both more participants and more data per participant. In fact, across the features in the sparse model, resampling analyses revealed that model contributions could only be reliably detected for the word onset feature frontally, and for lexical surprisal frontally and parietally (Fig 8B; note that frontal surprisal contributions are not shown, as these were only detectable with 40+ participants at 17+ min of data, at the lowest significance level of p < 0.05). Finally, while patterns of minimum data per participant required to reach significance from Figs 1–3 (i.e., when n = 41) are similar to those in Figs 7–8 (rightmost data points in each plot), in some cases the exact minutes per participants (or significance levels) don’t match between the two, since the true participant sample in Figs 1–3 corresponded to just one data point within the larger bootstrap distribution used for Figs 7–8.

**Figure 7.**
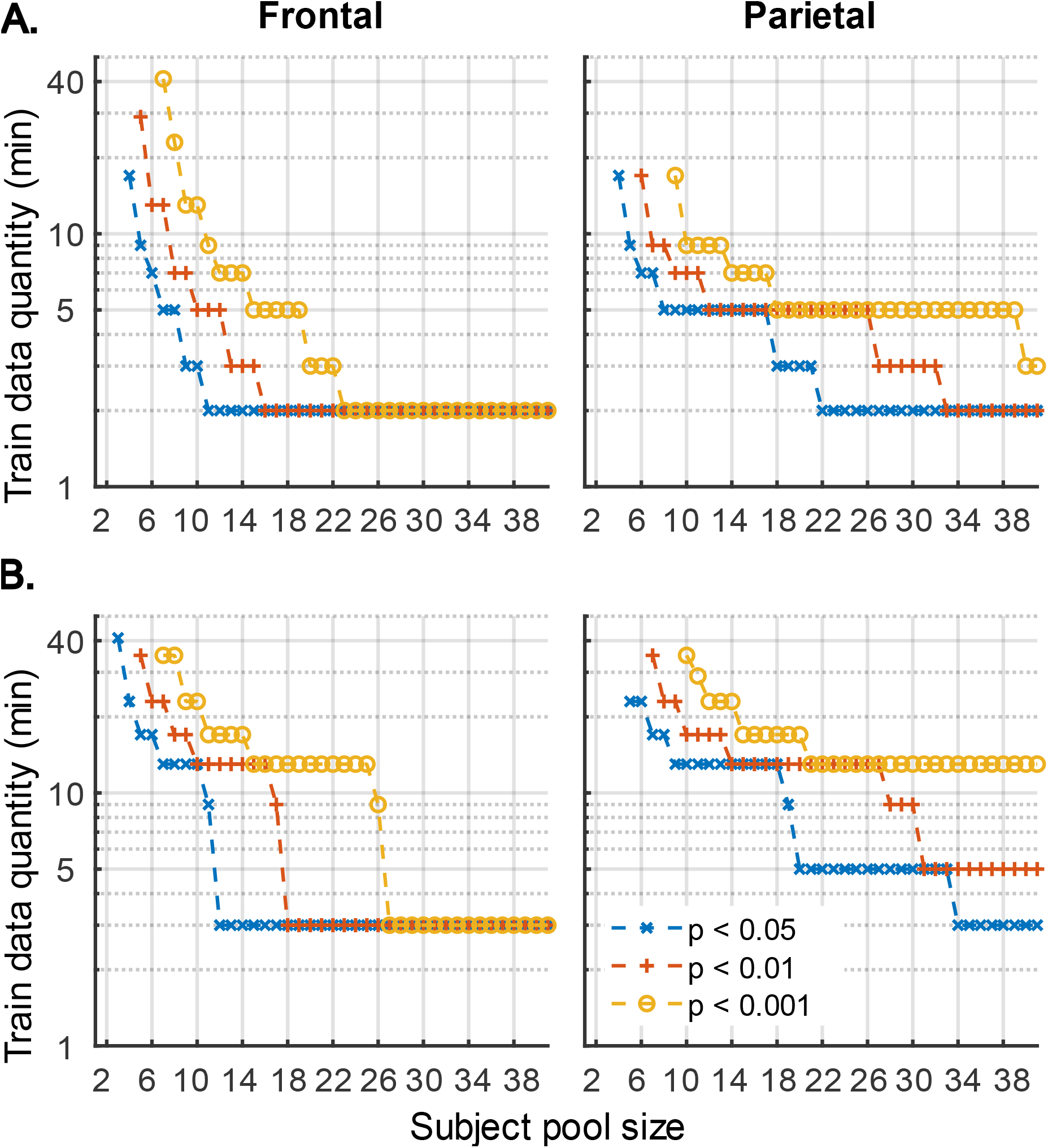
The amount of training data per participant required to reach significant overall prediction accuracy as a function of participant sample size for the denser (A) and sparser (B) TRF models in frontal (left column) and parietal (right column) ROIs. Different line colors represent different significance levels (see legend in lower right). For visualization purposes, minutes of data are shown on a log axis. The discrete steps along the y-axis stem from the fact that subject-specific analyses were run using 11 discrete data quantities per subject, the results of which were used in this resampling analysis. Therefore, values on the plots represent approximate threshold data quantities needed for each sample size.

**Figure 8.**
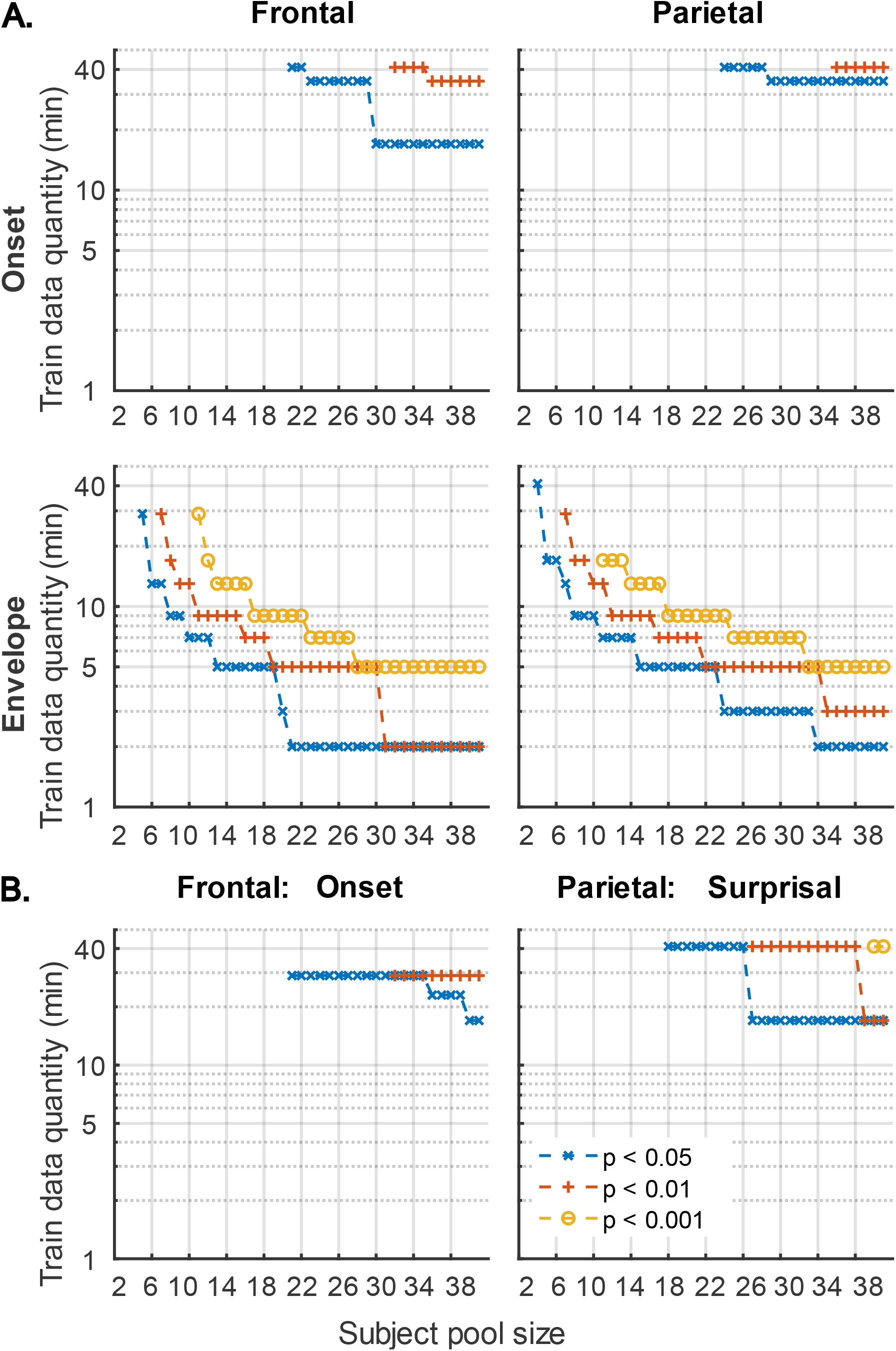
Amount of training data per participant required to reach significant feature-specific contributions to model prediction accuracies, as a function of participant sample size for the denser (A) and sparser (B) TRF models in frontal (left column) and parietal (right column) ROIs. Note that for the sparser model, only feature-ROI combinations where significant feature-specific model contributions could be achieved are shown (except for surprisal in the frontal ROI, where p < 0.05 could be achieved with 40+ participants at 17+ min of data). Line colors represent different significance levels (see legend in lower right).

### 3.2 Generic analyses

In the second set of analyses, we explored the effects of subject count on performance of *generic* analyses using the same two models (with denser and sparser features; section 2.6.3) used for subject-specific analyses (section 3.1). In contrast to subject-specific analyses, generic models are simultaneously fit to data from multiple participants and evaluated on their ability to predict data of held-out participants. Due to the higher computational load of fitting models to multiple subjects simultaneously, these models were tested on 2, 4 and 8 minutes of data per participant. Figures 9–14 depict results of generic analyses analogous to those shown in Figures 1–6 for subject-specific analyses. Note that the two sets of results are not directly comparable, as the latter results depict distributions over, and averages of, the central tendency of 20 resampled generic analyses, rather than distributions of individual subject results. The lower variability in these analyses is therefore not directly indicative of generic analyses performing better than subject-specific analyses. The resampling approach used here was important due to inherent noisiness and strong influence of outlier data in analyzing small subject counts (see section 2.6.2).

**Figure 9.**
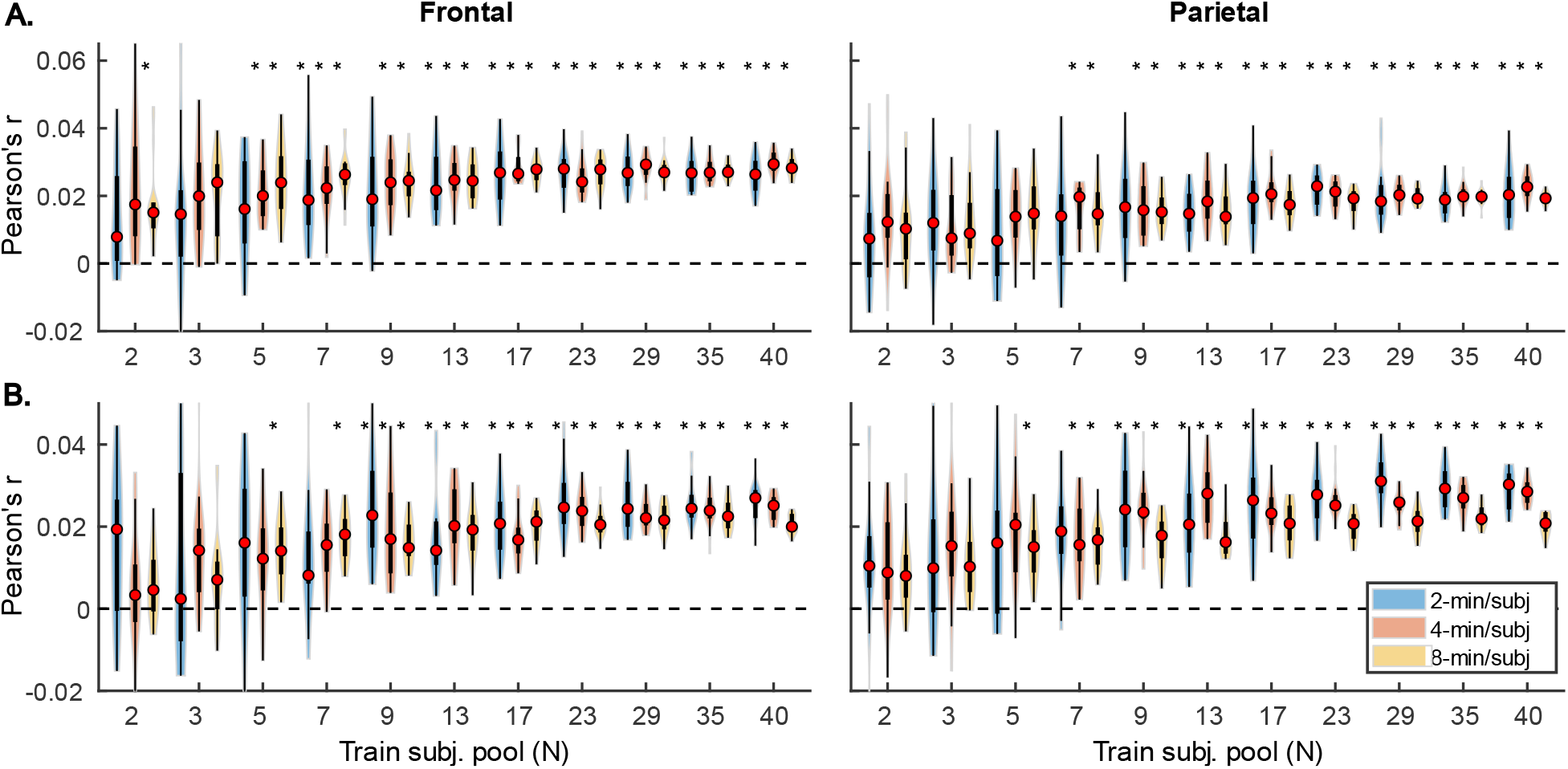
Noise floor-corrected generic model prediction accuracy distributions as a function of training subject pool size, for denser (A) and sparser (B) models with 2-min, 4-min, and 8-min of training data per subject (violin color; see legend in the lower left plot). The distributions depicted by violin plots represent collection of mean prediction accuracies across 20 resampling analyses (instead of across-subject variability seen in Figs 1–2). As such, only subject counts with distributions with no overlap with zero are deemed to be significant at p < 0.05 (uncorrected). Note that for each triplet of violins, the training subject pool is fixed with a value indicated by the x-tick label below the central violin of the triplet.

**Figure 10.**
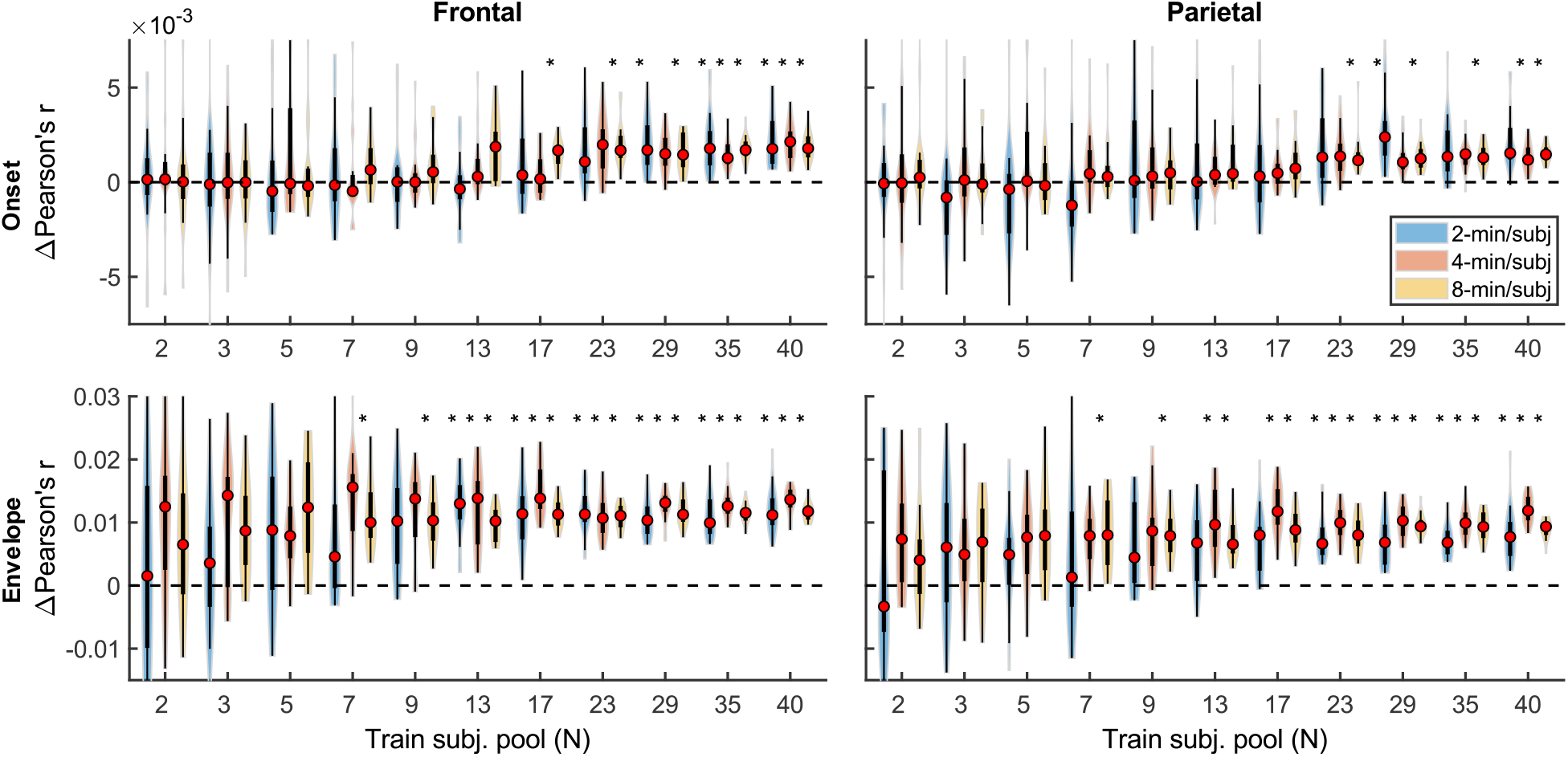
Feature-specific generic model contributions as a function of training subject pool size, for the denser model, with 2-min, 4-min, and 8-min of training data per subject (violin color; see legend in the upper left plot).

**Figure 11.**
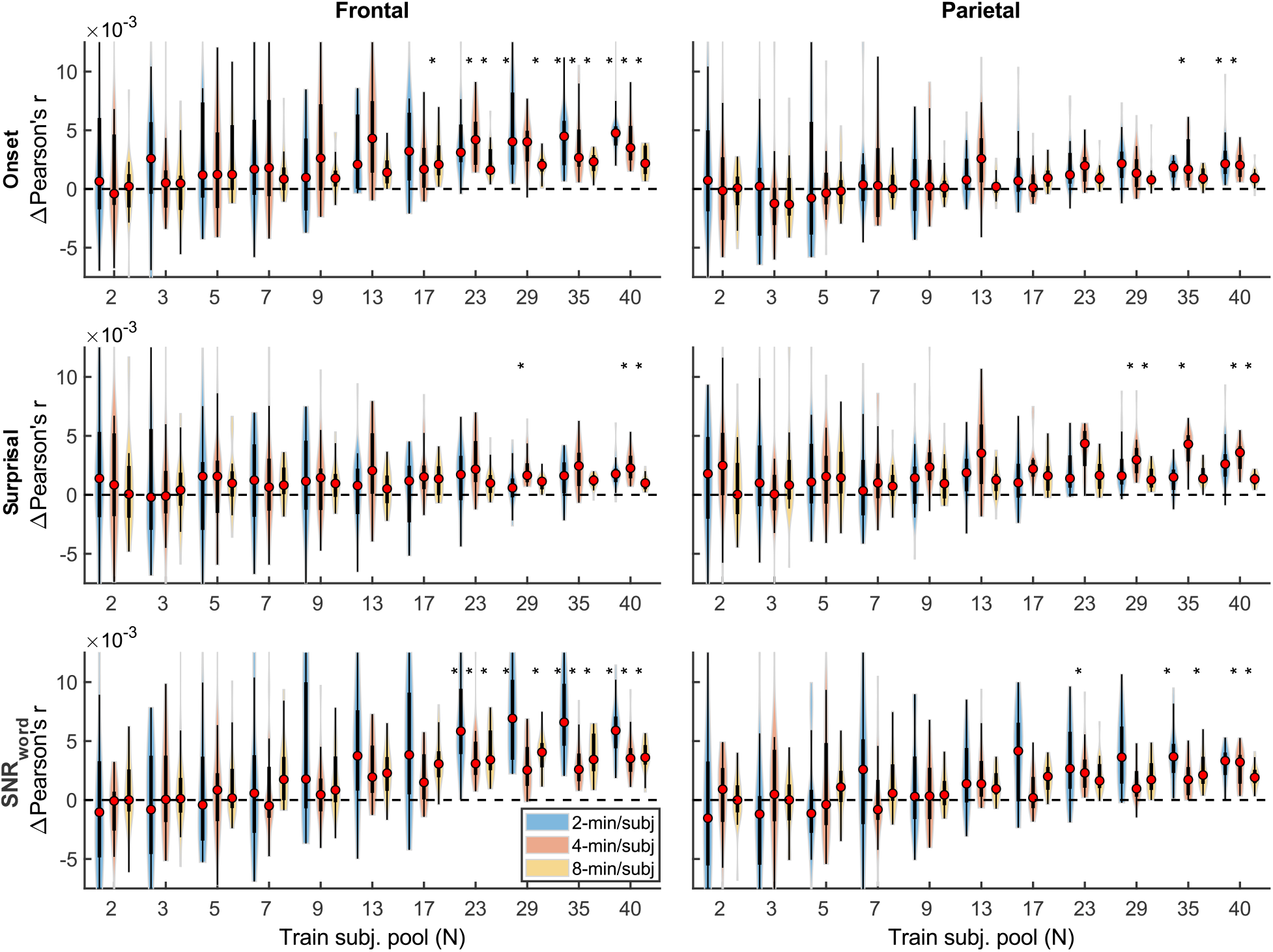
Feature-specific generic model contributions as a function of training subject pool size, for the sparser model, with 2-min, 4-min, and 8-min of training data per subject (violin color; see legend in the lower left plot).

**Figure 12.**
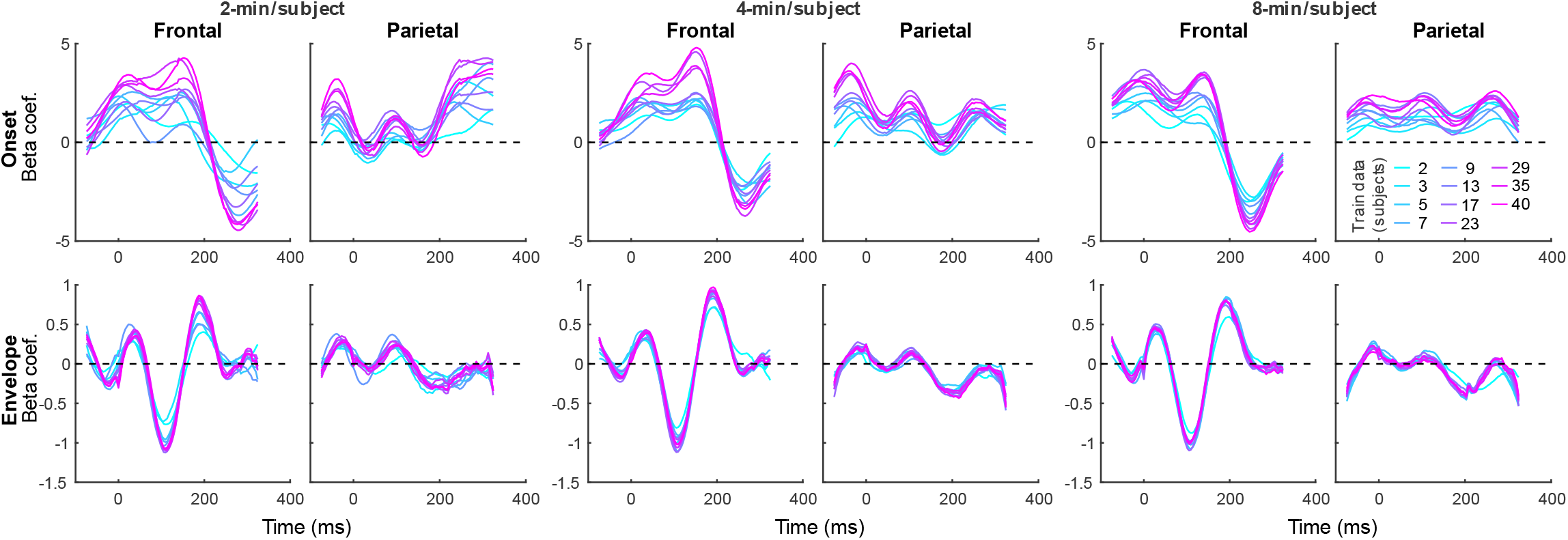
Average generic model TRF time courses for onset (top row) and envelope (bottom row) features from the denser model trained with 2-min (left pair of columns), 4-min (middle pair of columns), and 8-min (right pair of columns) of data per subject. TRFs estimated using different amounts of data are depicted in different colors (see legend in the upper right plot). Note that the sharp deflections in envelope response around t = 0 ms reflect mild leakage of electrical artifact from earphones into the EEG signal.

**Figure 13.**
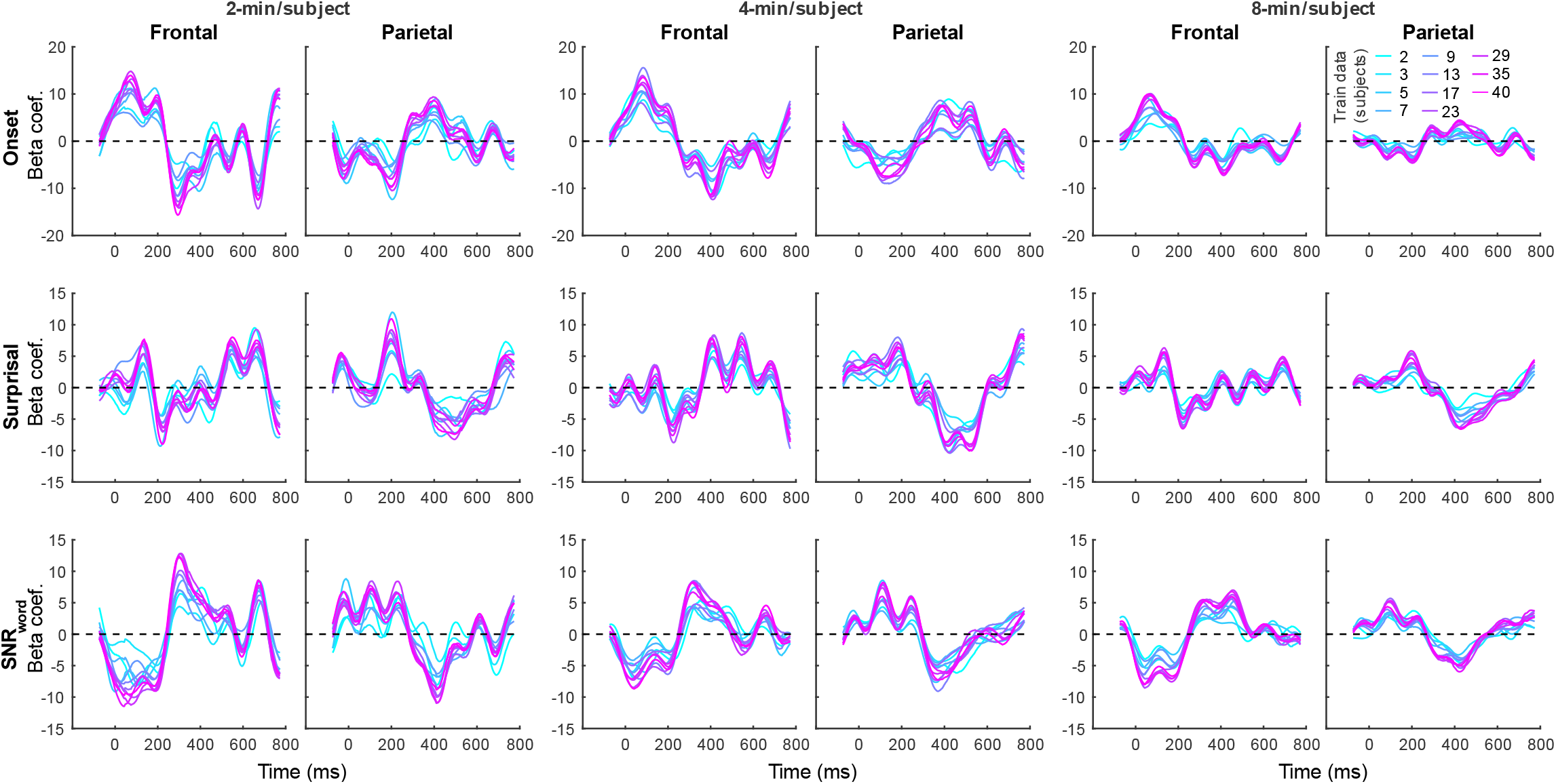
Average generic model TRF time courses for onset (top row), surprisal (middle row), and SNR_word_ (bottom row) features from the sparser model trained with 2-min (left pair of columns), 4-min (middle pair of columns), and 8-min (right pair of columns) of data per subject. TRFs estimated using different amounts of data are depicted in different colors (see legend above the upper right plot).

**Figure 14.**
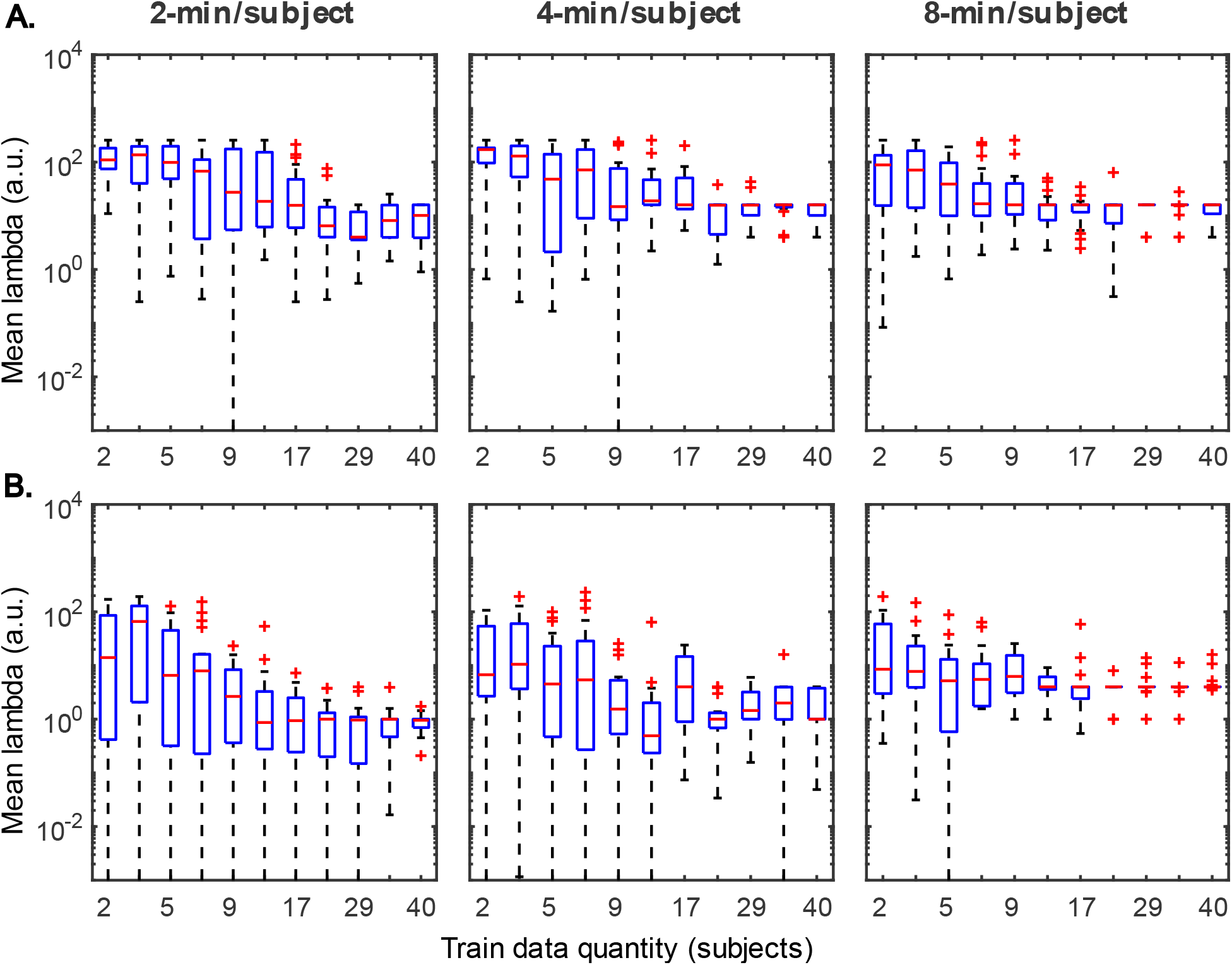
Distributions of generic model regularization parameters, lambda, from the denser (A) and sparser (B) models, as a function of training data quantity. Details of visualization are as in Fig 6.

Qualitatively, the generic model prediction accuracies (Fig 9) as well as model fit contributions (Figs 10–11) both exhibited apparent monotonically increasing performance and decreasing variability as the number of participants used to train the model increased. Because these plots depict distributions of average performances across 20 resampling analyses, only subject counts for which the entire distribution is elevated above zero can be deemed as reliably achieving significant non-zero performance. Quantitatively, however, bootstrap analyses of the linear relationship between the subject pool size used in model fitting and prediction accuracies (see section 2.7.2) only revealed a significant linear increase in overall prediction accuracies of the sparser model. None of the feature-specific model contributions in any ROI showed a significant relationship with the subject pool size. This is likely driven by the high variance of prediction accuracies with small numbers of participants. Nevertheless, it is evident from Figs 9–11 that model performance did improve with increasing participant pool, since significant prediction accuracies and feature-specific contributions were more consistently seen with increasing participant pools. Depending on the ROI, about 5-7 participants were needed to achieve reliably elevated overall model prediction accuracy, while substantially greater number of participants were needed to observe elevated feature-specific model contributions for different features in the denser and sparser models, respectively. A notable divergence between subject-specific and generic results was that in the former analyses, feature-specific contribution of surprisal but not SNR_word_ reached significance (Fig 3), whereas in the latter (Fig 11), SNR_word_ exhibited significant model contributions that were detected in a greater number of analyses than contributions of surprisal.

Interestingly, comparisons of 2-, 4- and 8-min of data per subject used in model fitting (different boxplot fill colors in each plot) made only a modest difference in performance, with the models trained on more data per participant generally producing tighter performance distributions, and reaching significance with fewer participants. However, it is noteworthy that the prediction accuracies and feature-specific contributions of the sparse model (Fig 9B, and Fig 11, respectively) exhibited a trend of decreasing peak prediction accuracies with increasing per-subject amount of training data (e.g., 8-min/subject models had slightly lower mean performance than 2-min/subject models, especially with larger participant pools). The cause of this is unclear and may warrant future exploration via analyses of different data (and feature) sets.

TRFs derived from denser generic models, averaged across the 20 resampling analyses, are depicted in Fig 12 (for easier comparison across data quantities, see heatmaps in Supplementary Figs S5-7). These TRFs were highly stereotypical across different subject pool sizes and the three per-subject data quantities (see supplementary Figs S11-S13 for correlation matrices of pairwise comparisons of TRFs). In line with subject-specific analyses, there was some evidence of increasing TRF amplitudes as the overall amount of data used to fit the model increased (i.e., with increasing number of subjects; see TRFs depicted by different colors in each subplot), although for envelope TRFs this effect diminished with increasing per-subject data quantities. This general patterns of increasing TRF amplitudes was again accompanied by a systematic decrease in the regularization parameters (Fig 14A). Notably, because most generic models were trained on greater total amount of data than any of the subject-specific models (for n > 20, n > 10, and n > 5, in models using 2-min, 4-min, and 8-min of data per subject, respectively), the upper bound of regularization parameter values seen in generic analyses was substantially lower than those seen in subject-specific analyses. These differences likely account for the slightly weaker envelope TRF amplitude increases as a function of training data quantity in generic analyses, compared to subject-specific analyses.

TRFs from the sparser model (Fig 13; Supplementary Figs S8-10) also exhibited a high degree of similarity for models trained on different amounts of data, although they generally show slightly higher degree of noisiness across different sample sizes than those derived using the denser model (except for dense model’s parietal onset TRFs; see supplementary Figs S14-S16). Mirroring the slight decrease in prediction accuracies with increasing per-subject data quantity (Fig 9B, Fig 11), there was an analogous decrease in TRF amplitudes (Fig 13, left vs right pairs of columns). At the same time, we also observed a pattern of slight increases in TRF amplitudes for models trained on more participants (i.e., different plot colors in each subplot of Fig 13; see heatmaps in Supplementary Figs S8-10), mirroring the decreasing amplitudes of regularization parameters in these analyses (Fig 14B). We speculate that this apparently paradoxical discrepancy between effects of more data per participant vs more participants could reflect tradeoff between phenomena driven by cognitive (e.g., waning attention or increased adaptation) vs technical (i.e., decreasing regularization parameter with more data) factors.

### 3.3 Similarity between subject-specific and generic TRFs

While subject-specific and generic analyses produced qualitatively similar TRFs (Figs 4 and 5 vs Figs 12 and 13), we sought to assess this similarity quantitatively using a correlation analysis (Fig 15). To this end, we focused on TRFs derived from the largest quantity of data in each analysis (including generic TRFs based on each of 2-min, 4-min and 8-min of data per subject), as these TRFs most accurately capture the underlying neural responses to each of the features. For each feature, we computed the Pearson’s correlation between TRFs derived in subject-specific and generic analyses. This analysis confirmed that in vast majority of cases, the two analyses produce highly similar TRFs, only seldom deviating below r = 0.75, predominantly for sparser features in ROIs with relatively low TRF amplitudes (e.g., parietal onset and frontal surprisal responses). These results demonstrate that at least for features modelled here, subject-specific and generic analyses enable extraction of highly similar neural responses.

**Figure 15.**
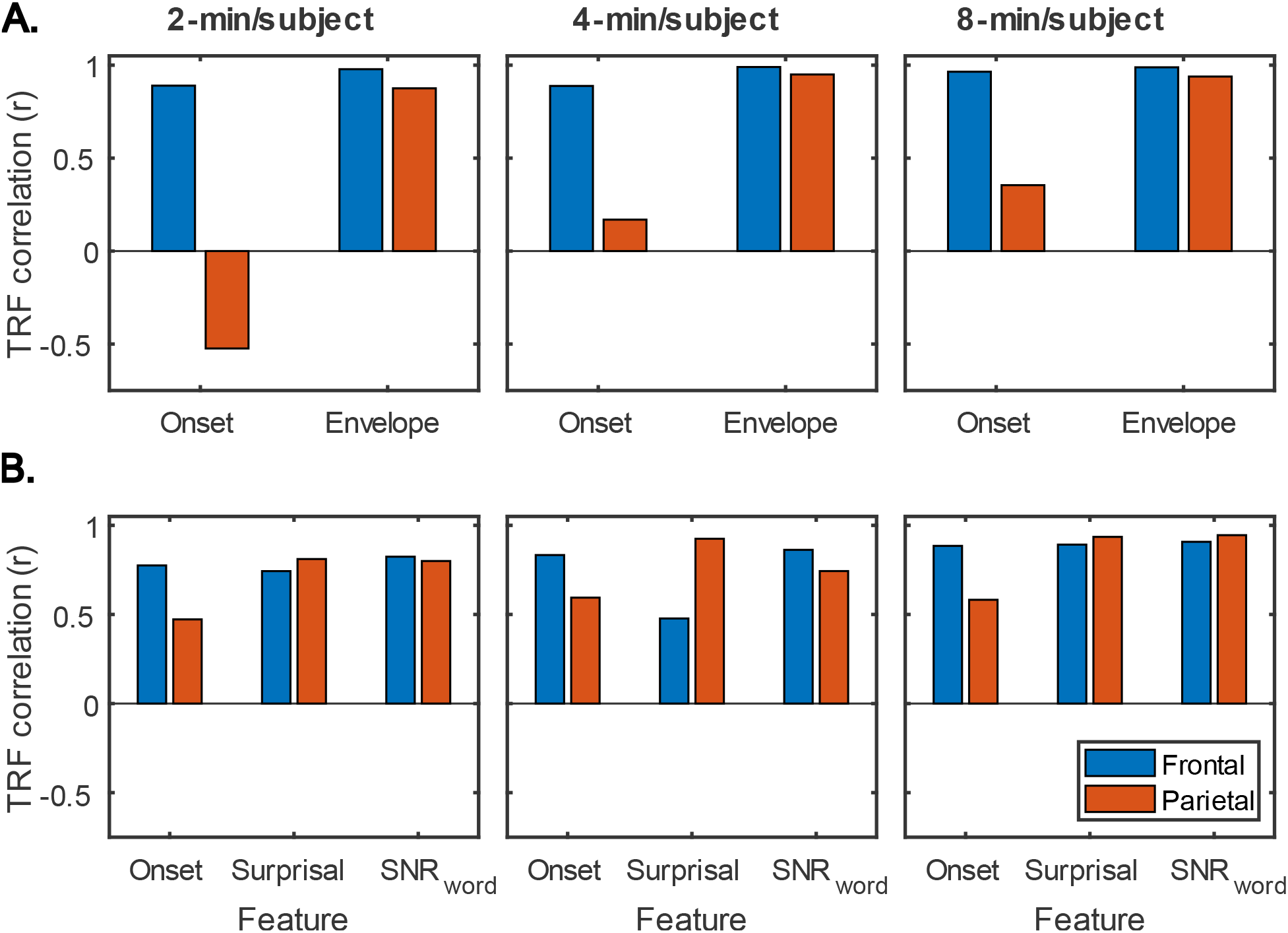
Similarity between average TRFs derived using subject-specific and generic analyses for features in the sparser (A) and denser (B) models. For generic analyses, both the results from analyses utilizing 2-min (left column), 4-min (middle column), and 8-min (right column) per subject are shown. Different bar colors represent the frontal and parietal ROIs (see legend in lower left plot). Note that these analyses utilized TRFs derived using models trained on largest amounts of data in each type of analysis, i.e., most data per subject for subject-specific analyses, and most subjects for generic analyses.

## 4. Discussion

### 4.1 Summary of goals and findings

TRF analyses (Lalor and Foxe, 2010; Crosse et al., 2016) of EEG and MEG data are increasingly popular in studies of cortical responses to continuous, naturalistic stimuli such as speech (e.g., Di Liberto et al., 2015; Broderick et al., 2018; Weissbart et al., 2019) and music (e.g., Di Liberto et al., 2020; Marion et al., 2021). However, relatively few informational and educational resources demonstrating the behavior of these analyses under various constraints exist. Such resources allow researchers who are new to TRF analyses or considering adopting them to gain key intuition and insight to guide their study design. The goal of the present work was to demonstrate how quantity of collected data, a key parameter in experimental design, influences TRF analyses of attended speech representations in the context of a dual-talker continuous speech paradigm. We addressed this question using a previously collected dataset (Mesik et al., 2021) using two types of analyses: 1) Subject-specific analyses in which TRF models are independently fit to each participant’s data, and 2) Generic analyses in which data from multiple participants is jointly used to fit a TRF model. For each analysis type we fit two different models, one of which had temporally denser features (acoustic envelope model), while the other had temporally sparser features (surprisal and SNR_word_ model). These models were fit repeatedly to explore how the amount of data per participant and number of participants influence model prediction accuracies in subject-specific and generic analyses, respectively. Finally, we used correlation analysis to compare the similarity of the TRFs derived in the two analysis approaches.

It is unsurprising that fitting models to more data resulted in monotonically improving prediction accuracies and more reliable TRF estimates. However, a closer examination of our results revealed several noteworthy phenomena. First, across both types of analyses, significant prediction accuracies could be achieved with just minutes of data per participant (Figs 1 and 9), although the denser model with envelope features had on average higher prediction accuracies than the sparser model with word-level features. Second, in analyses of feature-specific contributions to the overall prediction accuracies (Figs 2–3 and 10–11), we observed a marked dissociation between denser and sparser models. Specifically, feature-specific model contributions were generally much smaller for sparser features, and capturing these model contributions required much greater amount of training data. However, it is noteworthy that in generic analyses, even 2-4 minutes of data per participant were, in some feature-ROI combinations, enough to reveal these feature-specific model contributions (Fig 11). This demonstrates that signals related to sparse features may have sufficient signal-to-noise ratio to be detected even with relatively small amounts of data, provided that the model is trained on a sufficiently large dataset. Third, although TRFs derived with different amounts of data generally showed a high degree of time-domain consistency, we observed a systematic increase in TRF amplitudes with increasing data quantity (Figs 4–5, and 12–13). This pattern was mirrored by systematic decreases in the regularization parameter (Figs 6 and 14), which reflects the degree to which larger TRF amplitudes are penalized during model fitting. Finally, TRF patterns derived using subject-specific and generic analyses were highly similar (Fig 15), demonstrating that the two analyses reveal largely identical signatures of cortical speech processing.

### 4.2 Relationship to past literature

While most existing works utilizing TRF methods have used these tools to address specific questions about the nature of speech and music processing, only a handful of studies have explored the methodology itself. In general, the latter works focused on bigger picture overview of TRF methods and their utility in speech processing (Crosse et al., 2016; Sassenhagen, 2019), as well as on best practices in utilizing these methods in studies of special and clinical populations (Crosse et al., 2021). Additionally, Wong et al. (2018) explored performance of a range of regularization approaches for fitting forward and backward models in the context of a growing body of attention decoding literature.

Most closely related to the present work, Di Liberto and Lalor (2017) investigated the effect of data quantity on performance of subject-specific and generic forward models in the context of phoneme-level speech processing. Similar to our results, Di Liberto demonstrated that the ability of subject-specific models to capture responses related to the phonemic processing improved with greater amount of data, with models requiring about 30 minutes of data to do so reliably. Their generic model derived via averaging of subject-specific models was able to capture phonemic responses with 10-min of data, with no further improvement when more data per subject was used. While the prediction accuracies of our generic models improved with increasing subject pool, our results are consistent with those of Di Liberto in that we observed neither strong evidence of increases in generic model performance with increasing amount of data per participant, nor substantial increases in the mean feature-specific prediction accuracies with increasing subject pool. Nevertheless, utilizing larger training subject pools did result in decreased variance in the estimates of these contributions, and hence increased statistical power for their detection. Overall, our results agree with those of Di Liberto in that generic models can provide significant predictive power even when they are trained and evaluated on relatively small amounts of data per participant (2-/4-min and 10-min in our and Di Liberto’s studies, respectively).

### 4.3 Utility of subject-specific and generic TRF analyses

Given that generic analyses demonstrated superior performance to subject-specific analyses when small amounts of data per participant was available, it is important to consider the scenarios for which each analysis approach may be appropriate. Despite requiring more data per participant, subject-specific analyses have been overwhelmingly more popular than generic analyses in studies of speech and music processing. A key advantage of these analyses is that for each participant, subject-specific fits provide independent estimates of both prediction accuracies, and the TRFs themselves, allowing for traditional approaches to group-level statistics. Additionally, subject-specific modelling is critically important for studies seeking to characterize individual differences within a population, and/or their relationship to behavioral performance or other subject-level characteristics.

Conversely, while cross-validation used during generic model fitting also provides independent prediction accuracies for all participants, the TRFs from different cross-validation folds are non-independent. Moreover, the interpretation of prediction accuracies for individual subjects in the context of generic analyses differs from subject-specific approach, as they reflect the predictability of a given participant’s neural representations by a generic model, as opposed to the overall strength of speech representations in that participant. In other words, it may be the case that due to individual differences (e.g., due to anatomical variability) a particular participant’s data may be poorly predicted by a generic model even if their individual model could perform substantially better. However, by capturing the shared aspects of neural processing within a larger group, generic analyses may be particularly useful for categorizing participants, or their mental states, which may have important applications both in clinical diagnostics, and for practical tools such as neuro-steered hearing aid devices. Indeed, several studies utilizing *backward* models to decode attention have demonstrated the utility of generic models, albeit with a performance deficit relative to subject-specific models (e.g., Mirkovic et al., 2015; O’Sullivan et al., 2015). The potential of generic models for clinical applications is further supported by the key observation in the present work that they can achieve substantial predictive power on highly limited amounts of data per participant. However, as pointed out by Di Liberto and Lalor (2017), generic models implicitly assume within-group homogeneity in neural representations, which may be particularly questionable within clinical populations (e.g., Levy et al., 1997; Happé et al., 2006). As such, researchers need to be cognizant about this limitation. Finally, on a practical note, one disadvantage of generic analyses, as implemented in the present work, is that fitting them to large datasets requires large amounts of computational resources and time, especially when utilizing a resampling approach to fitting. However, efficient use of resources (e.g., downsampling and reducing data dimensionality via methods such as multiway canonical component analysis, MCCA; de Cheveigné et al., 2018), and alternative model estimation approaches, such as averaging of subject-specific models (e.g., Di Liberto & Lalor, 2017) or models fitted to subsets of participants, can mitigate these technical challenges.

Finally, it should be noted that in the broader context of their neuroscientific applications, TRF analyses have been utilized predominantly for discovery of scientific knowledge. However, due to their ability to extract complex neural responses to arbitrary stimulus features from M/EEG activity evoked by continuous, naturalistic stimuli, TRF analyses have potential for applications beyond basic science. As noted above, such applications may include clinical diagnostics of deficits in speech and music processing, as well as evaluation of interventions, such as hearing aids or sensory training. For example TRF analyses could aid in diagnosis speech perception deficits that are difficult to capture via traditional audiological tests (e.g., Tremblay et al., 2015), as well as perceptual consequences of cochlear synaptopathy (i.e., hidden hearing loss; see review by Liberman and Kujawa, 2017). Notably, backward models have recently started to show promise in revealing group-level effects of hearing aids on cortical representations of speech (e.g., Alickovic et al., 2020, 2021). However, while exploration of TRF methods for clinical uses in individual patients is an important future direction, significant methodological progress will be needed before such methods become viable in practice.

### 4.4 Limitations and methodological caveats

While the present work provides a detailed exploration of TRF model performance as a function of training data quantity, caution should be taken in generalizing our results to other TRF studies. Choices in the experimental design, data pre-processing, modelled feature space, and statistical analyses could all influence the performance and interpretation of TRF analyses (for a broader, more detailed overview of associated issues, see Crosse et al., 2021). Moreover, as TRF methods are growing in popularity, “best/recommended” analysis methods are likely to become outperformed as new and improved analysis and statistical methods emerge. Below we highlight a number of methodological choices made within this study that could have influenced our results, and briefly discuss alternative choices available to experimenters.

#### 4.4.1 Study design considerations

This work utilized a dataset obtained using a challenging dual-talker paradigm from a participant sample spanning a wide range of ages (18-70). Alternative stimulus choices, such as different talkers (e.g., male vs female; single vs multi-talker), types of noise (e.g., speech-shaped noise, multi-talker babble, etc.), spatial configuration of sound sources, and task can alter the overall prediction accuracies, and TRF amplitudes. For example, neural representations of unattended speech are generally substantially weaker (e.g., Ding and Simon, 2012; Mesgarani and Chang, 2012), or even absent in the case of higher-order features related to phonetic (Teoh et al., 2022) and semantic processing (Broderick et al., 2018). It is notable that most existing speech TRF studies utilized single-talker paradigms. It is likely that for a model with fixed complexity, these paradigms require appreciably lower data quantities to achieve significant prediction accuracies, given their high speech intelligibility. Indeed, in the context of subject-specific models, several single-talker works have reported significant feature-specific contributions for higher-order features with as little as 12 minutes of data (e.g., Broderick et al., 2021; Gillis et al., 2022b). We chose to use a dual-talker paradigm in part because studies exploring neural correlates of speech perception difficulties are increasingly popular, making the present analyses more relevant to these investigators. Finally, despite using a dual-talker paradigm, it is likely that the TRFs obtained here are representative of TRFs one may derive in single-talker paradigms, as similar TRFs have previously been observed using both paradigms (e.g., Kong et al., 2014; Broderick et al., 2018).

In terms of participant sample, prior work has shown amplified cortical tracking of speech in older (e.g., Presacco et al., 2016; Decruy et al., 2019; Zan et al., 2020; Mesik et al., 2021, but see Gillis et al., 2022b) and hearing-impaired populations (Decruy et al., 2020; Fuglsang et al., 2020; Gillis et al., 2022a). As such, the inclusion of older participants in our data set may have resulted in slightly higher prediction accuracies and hence a need of somewhat less data to reach a criterion level of model performance, relative to what one may expect from data obtained only from younger adults. Note that we chose to utilize the combined data set given that in our previous work (Mesik et al., 2021) younger and older adults exhibited similar temporal morphologies of word-level TRFs, and hence it is likely that these groups’ speech-driven responses vary along a continuum, rather than reflecting categorically different speech processing (the latter of which would make combined analysis of the two sets of participants invalid). Lastly, we caution the reader that the present results are likely less applicable to special populations, such as clinical patients or children, whose speech processing may be immature or otherwise altered relative to the healthy adult population studied here.

#### 4.4.2 Pre-processing considerations

Pre-processing operations applied to data in order to improve its SNR, likewise, can have substantial impact on TRF analyses. Filtering is one of the most common pre-processing operations, allowing researchers to isolate a subset of frequencies thought to contain the neural signals of interest, while removing contributions of noise sources. While some filtering is usually desirable, we caution the reader that different types of filters can introduce systematic distortions into the data, which can negatively impact the interpretability of the extracted neural signals, such as ERPs or TRFs. We refer the reader to a detailed overview of issues related to filtering by de Cheveigné and Nelken (2019). In the present work, data was filtered to isolate the neural dynamics in the low-frequency delta (1-4 Hz) and theta (4-8 Hz) bands of cortical responses to speech. Although this was largely motivated by previous findings that speech processing mechanisms track predominantly the low-frequency dynamics of speech signals (e.g., Zion Golumbic et al., 2013), auditory cortex is known to phase-lock to acoustics features at rates up to 100 Hz (e.g., Holmes et al., 2018). Thus, depending on the nature of studied feature representations, different filter cutoffs than those used here may be required.

Besides temporal filtering, of particular interest to TRF analyses are more sophisticated denoising algorithms, such as the denoised source separation (DSS; de Cheveigné and Simon, 2008), and multiway canonical correlation analysis (MCCA; de Cheveigné et al., 2018). DSS is a dimensionality reduction technique that can isolate components of neural activity most strongly related to a particular stimulus representation, such as the acoustic envelope, thus providing improved SNR for subsequent TRF analyses. This technique may be used to improve SNR in subject-specific analyses, likely achieving significant predictive power with lower amounts of data. Similarly, MCCA is a technique that isolates components of neural activity shared across subjects within a data set, which allows researchers to factor out subject-level sources of variability, such as differences in anatomy. MCCA, therefore, is particularly relevant for generic analyses that fit models to multiple participants simultaneously. In present work, generic analyses assumed anatomical alignment of neural activity at the level of EEG sensors. Differences in anatomy may have therefore contributed to lower prediction accuracies than could likely be achieved with MCCA.

#### 4.4.3 Modelled feature spaces

One of the key aspects of TRF analysis is the choice of the modelled feature space. Here we fit two separate models emphasizing denser and sparser features in order to compare the relative performance of these types of feature sets. However, it is important to highlight that this separation of features into distinct models served a demonstrative role, rather than being methodologically necessary (but see discussion of banded ridge regression in section 4.4.4). Fitting of models with smaller subsets of features can be particularly problematic when the scientific question concerns whether a given feature appears to explain unique variance in cortical responses. Specifically, it may be the case that variance explained by one feature can be explained similarly or better by other, correlated features that were not included in the model. By using a more complete feature representation, researchers can gain greater (albeit not complete) certainty about the unique explanatory power of different features in the model. Indeed, several recent studies have brought into question unique cortical tracking of higher-order features, such as phonemic categories (Daube et al., 2019; but see Teoh et al., 2022) and semantic dissimilarity (Gillis et al., 2021), when more complete representations of stimulus acoustics were included in the model. In this regard, we acknowledge that the present work does not conclusively establish that the modelled features per se were uniquely tracked in our data.

Finally, while most existing TRF studies have modelled responses to researcher-defined stimulus features, a notable emerging approach relies on passing stimuli through pre-trained artificial neural networks (e.g., deep neural networks; DNNs), and deriving feature sets from the abstract representations that emerge within these networks. This approach has been successfully used in functional magnetic resonance imaging (fMRI) studies (e.g., Güçlü and van Gerven, 2015; Kell et al., 2018), and given its promise, it will likely become increasingly popular in TRF analyses as well. However, while fitting more complex models can generally account for greater amount of variance, they introduce several challenges that researches need to deal with. First, a practical issue is that larger models require larger amounts of data, in order to more thoroughly sample the feature space included in the model. As such, the quantities of data that were sufficient to fit our relatively small models may be insufficient for adequate fitting of substantially larger models, such as ones with feature spaces based on activation patterns of DNNs. Second, uninformative features may be particularly common in large models, which can lead to overfitting of the training data set and poorer test performance. Pruning (i.e., systematic elimination) of uninformative features is therefore especially important with these larger models. Finally, while larger models can more accurately predict brain responses, they also tend to be more challenging to interpret, particularly when their features are derived using artificial neural networks. A major challenge in the interpretation of larger models is determining what cognitive processes (or neural computations) does tracking of various abstract features reflect.

#### 4.4.4 TRF analysis methods

The TRF estimation methods themselves offer researchers several options that may affect the performance of TRF analyses. In the present work, we fit TRF models using regularized linear regression, while other existing methods include boosting (David et al., 2007; for boosting-based toolbox for M/EEG analyses, see Brodbeck et al., 2021), normalized reverse correlation (Theunissen et al., 2001), as well as a method for TRF estimation in the neural source space (Das et al., 2020). Although these alternative methods are beyond the scope of the present work, we note that Kulasingham and Simon (2022) showed that with a fixed amount of training data, regularized linear regression and boosting algorithms result in highly similar TRF estimates, making it likely that most of the patterns observed in the present work would also apply to boosting-based analyses. However, since boosting does not utilize a regularization parameter, it is unclear whether similar dependence of TRF amplitudes on training data quantity would be observed with boosting.

With respect to regularization methods used in the present work, it is notable that each analysis utilized a single ridge parameter, meaning that the fitting procedure penalized regression (beta) coefficients of all features included in the model to the same extent. However, because different features may have substantially different predictive powers, and may correlate with one another to different extents, single regularization parameter may be insufficient to allow for accurate TRF estimation of all features in a model (Nunez-Elizalde et al., 2019). To account for this issue, banded ridge regression methods have recently been introduced, which apply individualized ridge parameters to different features in a model (Nunez-Elizalde et al., 2019; Dupré la Tour et al., 2022). This method was not used in the present work, as we chose to build our analysis pipelines using publicly available tools (i.e., mTRF toolbox), which did not have this functionality implemented, yet (but see Crosse et al., 2021 for the description of forthcoming banded ridge regression functionality). Given our use of relatively small feature sets, it is unlikely that banded ridge regression would lead to substantial improvements in the performance of our analyses. Nevertheless, exploration of the effects of training data quantity on banded ridge regression performance is a useful future direction.

Estimation of feature-specific model contributions is another aspect of TRF analysis with multiple existing methods. Here we utilized perhaps the most common approach, which computes feature-specific contributions as the difference in prediction accuracies between full and reduced models containing all but one feature (e.g., Di Liberto et al., 2015; Gillis et al., 2021). Another approach utilizes comparisons of full and “null” models with equal dimensionality, where regressors of individual features are shuffled to disrupt their temporal relationship with the stimulus (e.g., Brodbeck et al., 2018). In certain cases, performances of these types of null models have been computed from full model’s TRF estimates (Broderick et al., 2021; Mesik et al., 2021), although potential biases this may introduce in fit contribution estimates have not been thoroughly assessed. The partial correlation approach for estimating feature-specific model contributions relies on comparisons of predictions of models utilizing individual feature representations (e.g., Prinsloo and Lalor, 2022; Teoh et al., 2022). Lastly, feature-specific contributions can also be estimated by subtracting (i.e., partialling out) predictions of an all-but-one feature model from the EEG data, and then fitting this residual signal with a model containing the left-out feature (O’Sullivan et al., 2021). In general, we anticipate that most of these methods result in correlated estimates of model contributions, albeit more work will be needed to assess their relative performances.

Finally, we reiterate that the ROI-based statistical analyses in the present work were used partly to highlight general locations where the two models performed well, and partly to simplify the presentation of results. As such, it is possible that different subsets of electrodes, identified using, e.g., cluster-based permutation tests (Maris and Oostenveld, 2007), could be more sensitive for detection of significant prediction accuracies and feature-specific model contributions. Use of these methods could, likewise, lead to increased statistical power, and consequently lower required training data quantity.

### 4.5 Conclusions

The goal of this work was to develop an informational resource for the growing field of TRF analyses of continuous speech processing, demonstrating the behavior of TRF analyses as a function of data quantity used in TRF fitting. In the context of relatively simple models of lower-level envelope processing, as well as higher-order processing of word-level features, we demonstrate that given a large-enough participant pool, small amounts of data (< 5 min) can be sufficient to train subject-specific models that predict significant variance in EEG responses to speech-masked speech. At the same time, substantially more data (15+ min) may be needed to capture feature-specific model contributions of individual word-level features. On the other hand, generic models can support significant prediction accuracy even for feature-specific variance with as little as 2-min of data per participant, while providing highly similar TRF estimates to those seen in subject-specific analyses. As such, despite their infrequent use, generic models have potential to be particularly useful for applications in clinical diagnostics, and multi-task studies with low per-task time budgets. While the present work is not, on its own, intended to be prescriptive about experimental duration, it may be a useful resource for informing selection of experimental duration, especially in conjunction with other tools, such as simulations and piloting.

## Supporting information

All supplemental materials

## Data availability

Data and analysis scripts will be made available upon reasonable request. Requests should be directed to JM, or MW.

## Ethics statement

The Institutional Review Board of the University of Minnesota approved the procedures in this study. All participants provided written informed consent to participate.

## Author contributions

JM and MW designed the original experiment, analyzed the data, and wrote the manuscript. JM implemented experimental procedures and collected the data. All authors commented on the manuscript and approved the submitted version.

## Acknowledgments

We thank Lucia Ray for help with carrying out the original study that yielded the data used in the present work.

