## Supplementary material for "The effects of data quantity on performance of temporal response function analyses of natural speech processing": All supplemental materials

A. Denser model

| | df | F | p |
|---|---|---|---|
| (Intercept) | 1, 47.5 | 120.1 | < 0.001 |
| datQnt | 1, 861 | 135.9 | < 0.001 |
| ROI | 1, 861 | 69.0 | < 0.001 |
| datQnt*ROI | 1, 861 | 4.1 | 0.042 |

B. Sparser model

| | df | F | p |
|---|---|---|---|
| (Intercept) | 1, 50.1 | 34.1 | < 0.001 |
| datQnt | 1, 861 | 163.5 | < 0.001 |
| ROI | 1, 861 | 10.0 | 0.002 |
| datQnt*ROI | 1, 861 | 2.1 | 0.150 |

Supplementary Table 1. LMM ANOVA for effects of data quantity and ROI on overall prediction accuracy in the denser (A) and sparser (B) models. Data quantity is represented by "datQnt." In both models, highly significant effect of data quantity was observed, reflecting the increases in prediction accuracy as a function of increasing data quantity.

A. Denser model

| Onset | df | F | p |
|---|---|---|---|
| (Intercept) | 1, 68.8 | 0.3 | 0.588 |
| datQnt | 1, 861 | 13.1 | < 0.001 |
| ROI | 1, 861 | 0.4 | 0.514 |
| datQnt*ROI | 1, 861 | 0.1 | 0.743 |

| Envelope | df | F | p |
|---|---|---|---|
| (Intercept) | 1, 54 | 87.7 | < 0.001 |
| datQnt | 1, 861 | 13.2 | < 0.001 |
| ROI | 1, 861 | 8.2 | 0.004 |
| datQnt*ROI | 1, 861 | 1.4 | 0.240 |

B. Sparser model

| Onset | df | F | p |
|---|---|---|---|
| (Intercept) | 1, 78.8 | 0.0 | 0.874 |
| datQnt | 1, 861 | 3.1 | 0.080 |
| ROI | 1, 861 | 0.3 | 0.607 |
| datQnt*ROI | 1, 861 | 4.7 | 0.031 |

| Surprisal | df | F | p |
|---|---|---|---|
| (Intercept) | 1, 82.4 | 2.2 | 0.139 |
| datQnt | 1, 861 | 0.8 | 0.375 |
| ROI | 1, 861 | 0.5 | 0.474 |
| datQnt*ROI | 1, 861 | 1.7 | 0.199 |

| Word_SNR | df | F | p |
|---|---|---|---|
| (Intercept) | 1, 72.1 | 2.0 | 0.162 |
| datQnt | 1, 861 | 8.3 | 0.004 |
| ROI | 1, 861 | 17.4 | < 0.001 |
| datQnt*ROI | 1, 861 | 21.0 | < 0.001 |

Supplementary Table 2. 1. LMM ANOVA for effects of data quantity and ROI on feature-specific model contributions in the denser (A) and sparser (B) models. The upper left cell of each table indicates the feature that ANOVA results correspond in that table corresponds to. While data quantity effects were robust in the denser model, they were only apparent for $word_{SNR}$ in the sparser model. However, given that $word_{SNR}$ did not show reliably elevated contributions even at large data quantities, we hesitate to interpret the significant effect of data quantity from its LMM ANOVA.

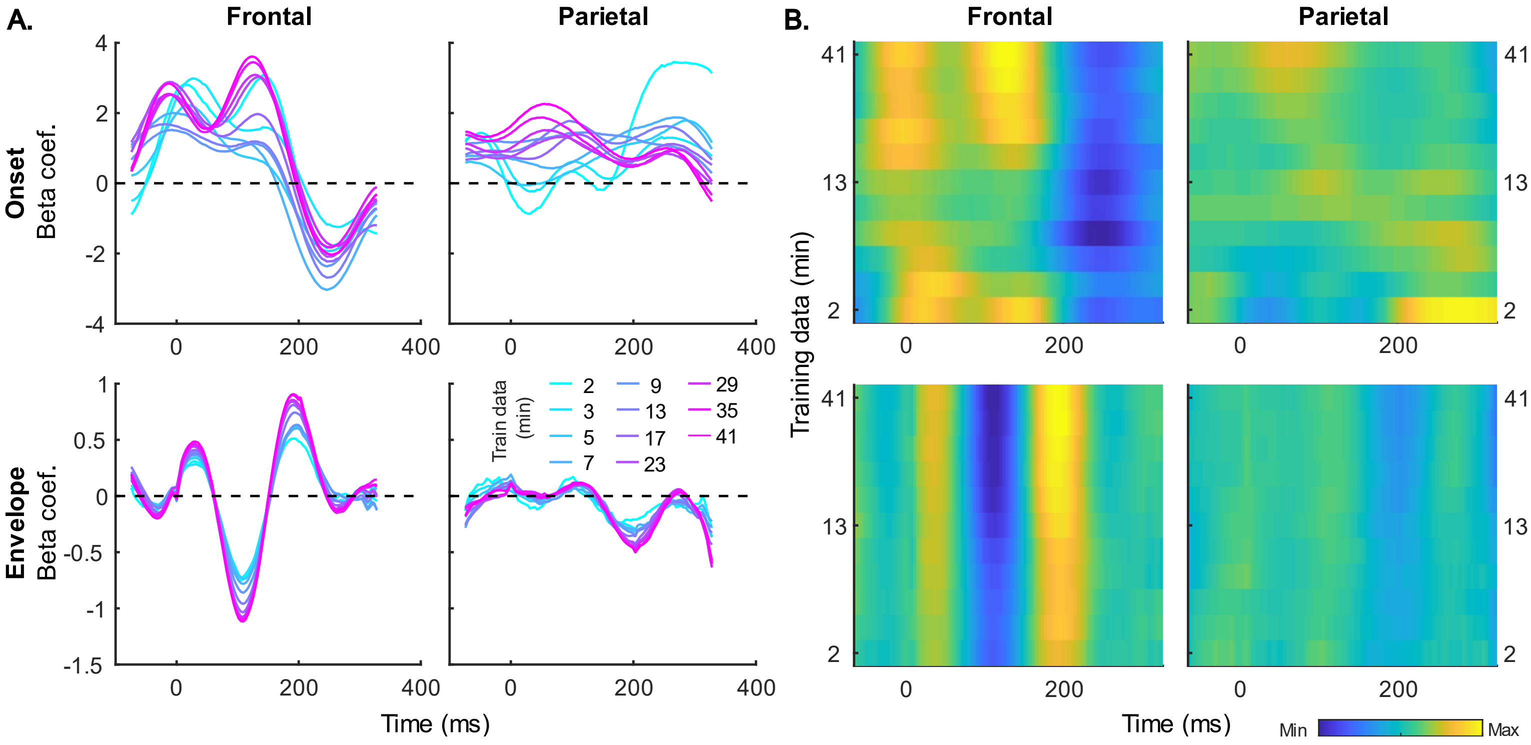


Supplementary Figure S1. Group-averaged TRF time courses of model beta coefficients for onset (top row) and envelope (bottom row) features from the denser model, depicted in line (A) and heatmap (B) forms. TRFs from frontal and parietal ROIs are shown in the left and right columns of each plot type, respectively. In line plots, increasing data quantity is reflected in line color (see legend in the lower left plot), while TRFs derived using higher quantity of data are in higher rows of the heatmaps. Note that heatmap plots are included to allow for easier comparison of relative TRF amplitudes as a function of data quantity.

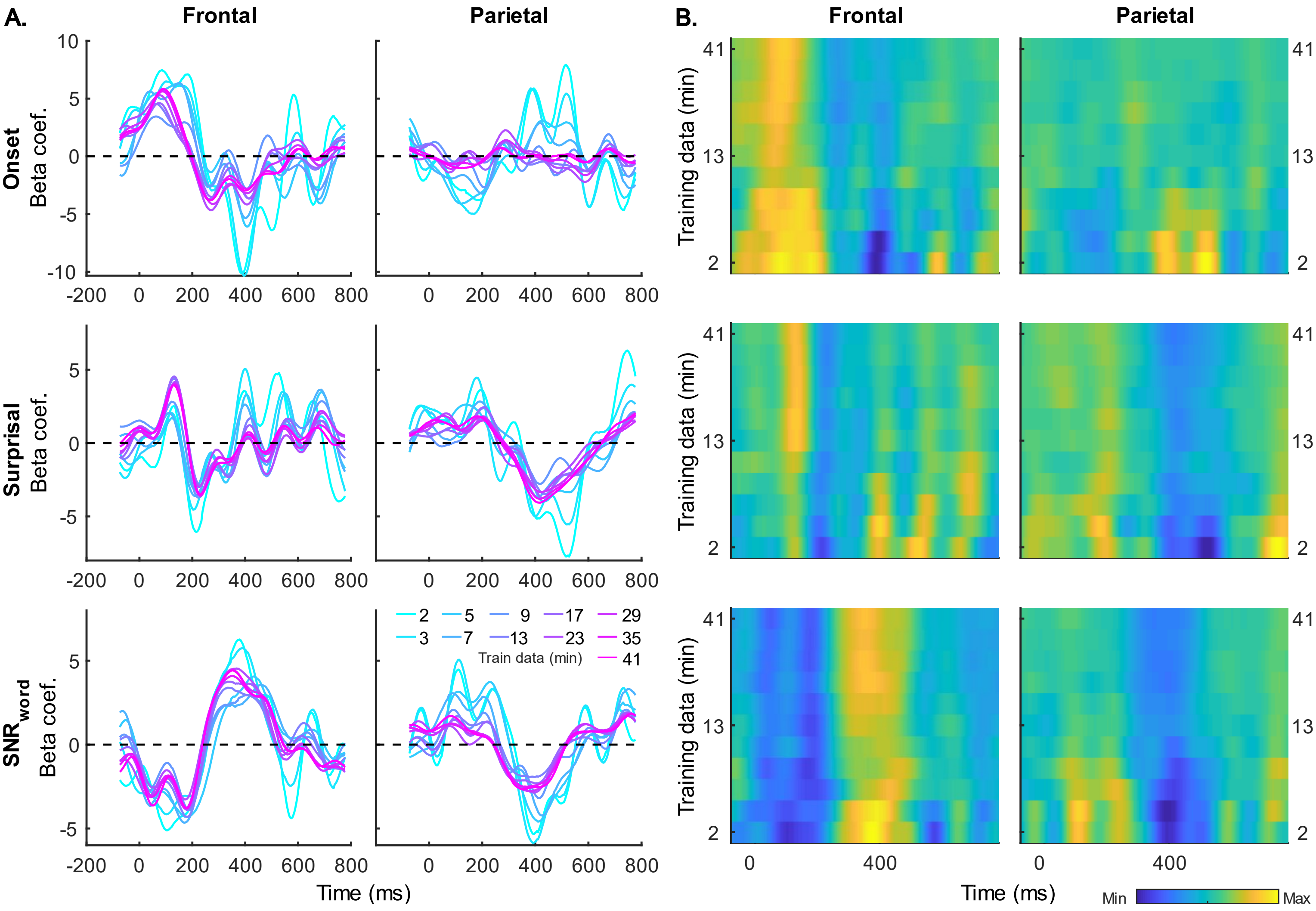


Supplementary Figure S2. Group-averaged TRF time courses of model beta coefficients for onset (top row), surprisal (middle row), and $SNR_{word}$ (bottom row) features from the sparser model, depicted in line (A) and heatmap (B) forms. Plotting conventions are as in Fig S1.

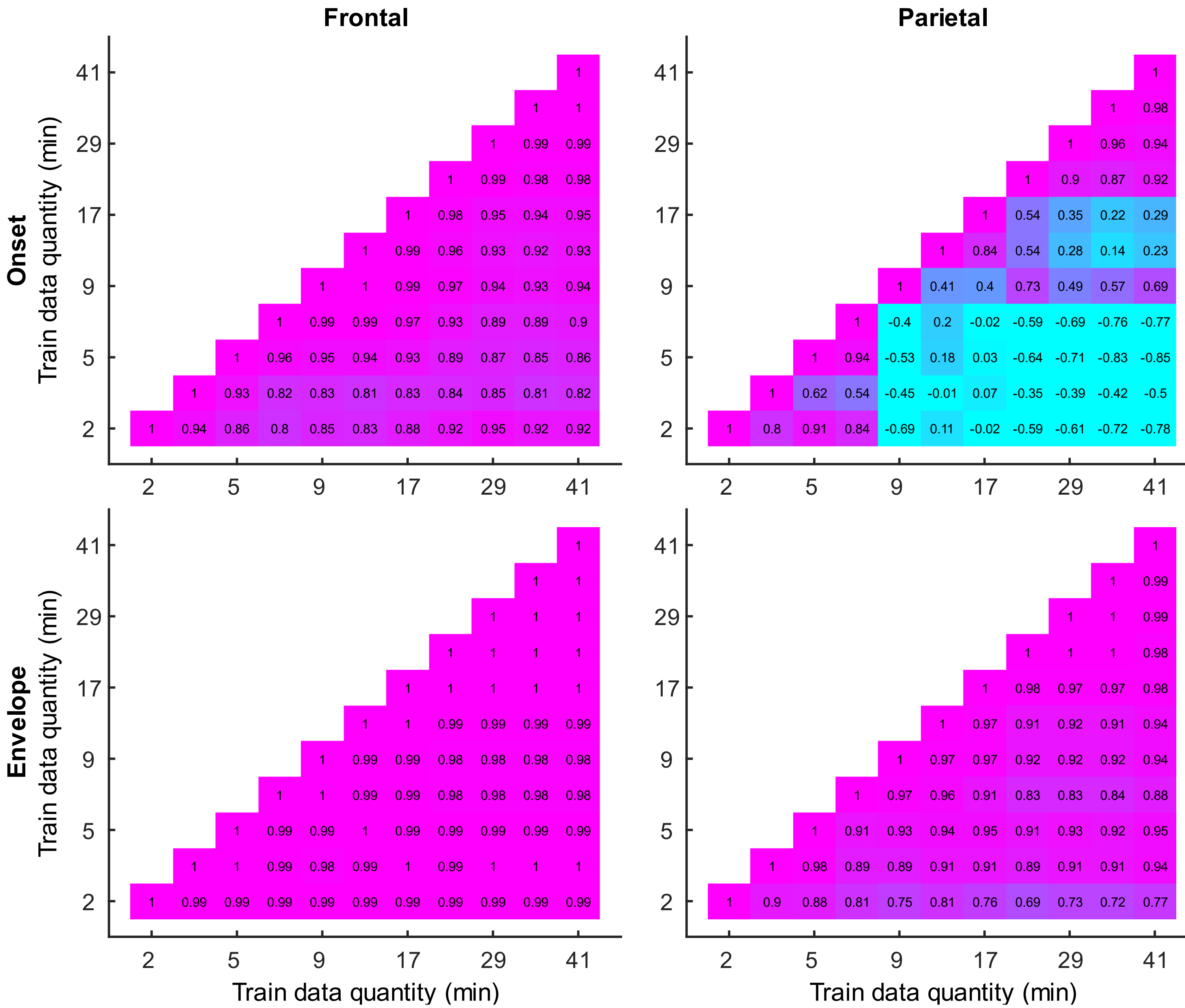


Supplementary Figure S3. Similarity of denser model's TRFs derived using different amounts of data for onset (top row) and envelope (bottom row) features in frontal (left column) and parietal (right column) ROIs. Each plot depicts pairwise Pearson's correlations between TRF time courses for all combinations of training data quantities (indicated along x and y axes of the plots). Brighter purple and blue colors reflect higher positive and negative correlation coefficients, respectively. Coefficient values are also indicated numerically at the center of each cell.

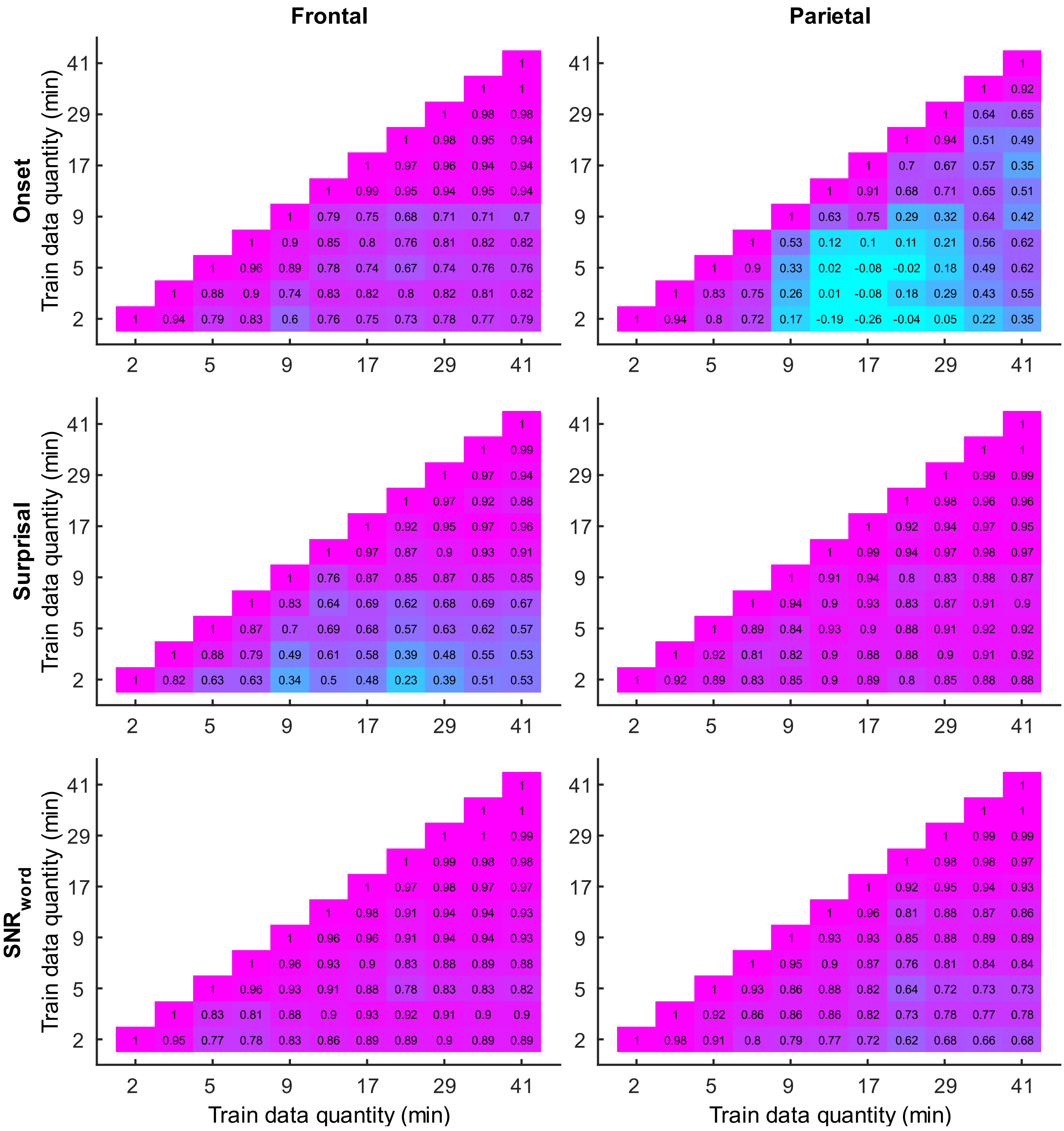


Supplementary Figure S4. Similarity of sparser model's TRFs derived using different amounts of data for onset (top row), surprisal (middle row), and $SNR_{word}$ (bottom row) features in frontal (left column) and parietal (right column) ROIs. Visualization conventions are as in Fig S3. Note the general trend of lower correlations for these features, compared to those in the denser model (Fig S3).

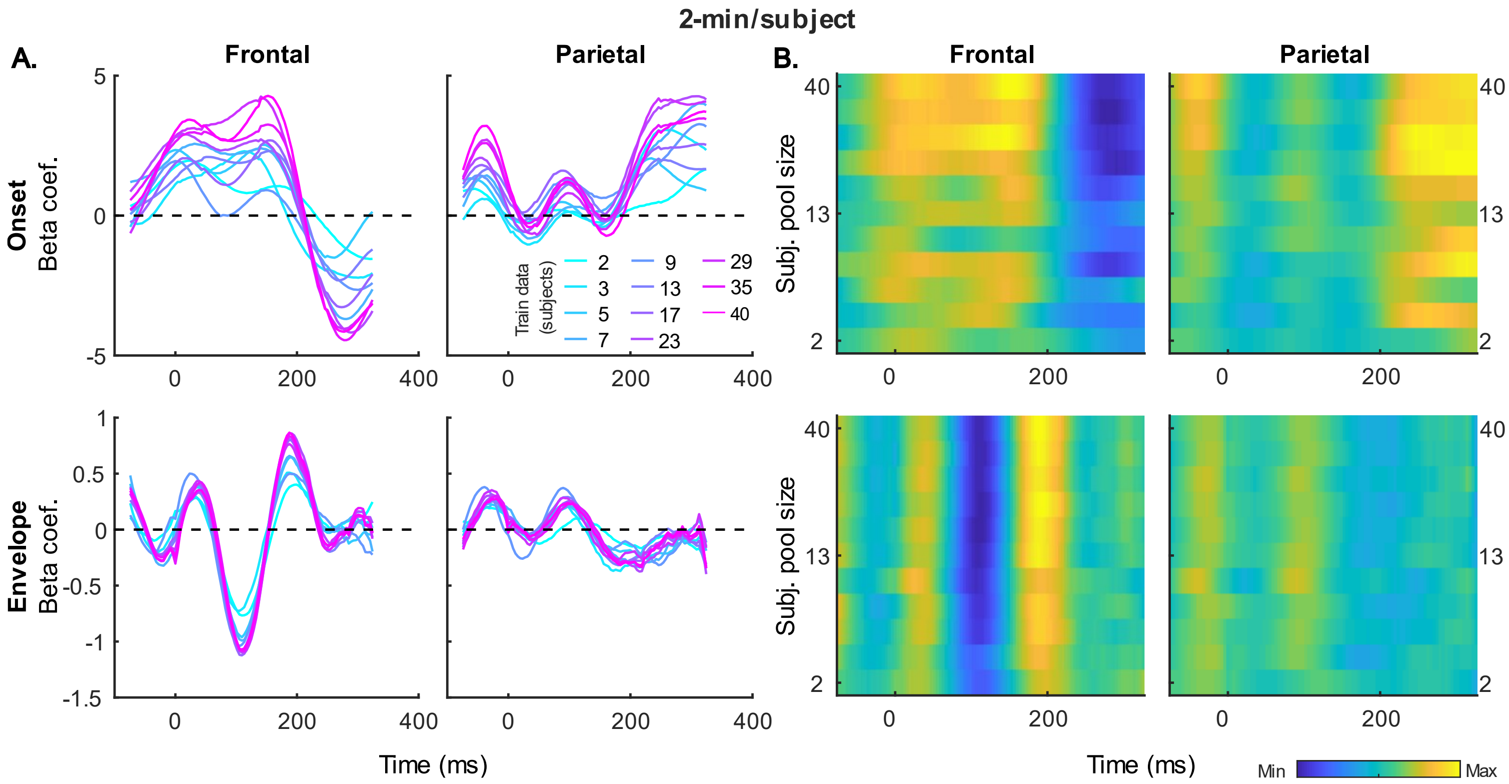


Supplementary Figure S5. Generic model TRF time courses, averaged across 20 resampling analyses, of model beta coefficients for onset (top row) and envelope (bottom row) features from the denser model trained with 2-min of data per participant. Same TRFs are shown in the form of line (A) and heatmap (B) graphs. TRFs from frontal and parietal ROIs are shown in the left and right columns of each plot type, respectively. In line plots, increasing data quantity is reflected in line color (see legend in the lower left plot), while TRFs derived using higher quantity of data are in higher rows of the heatmaps. Note that heatmap plots are included to allow for easier comparison of relative TRF amplitudes as a function of data quantity.

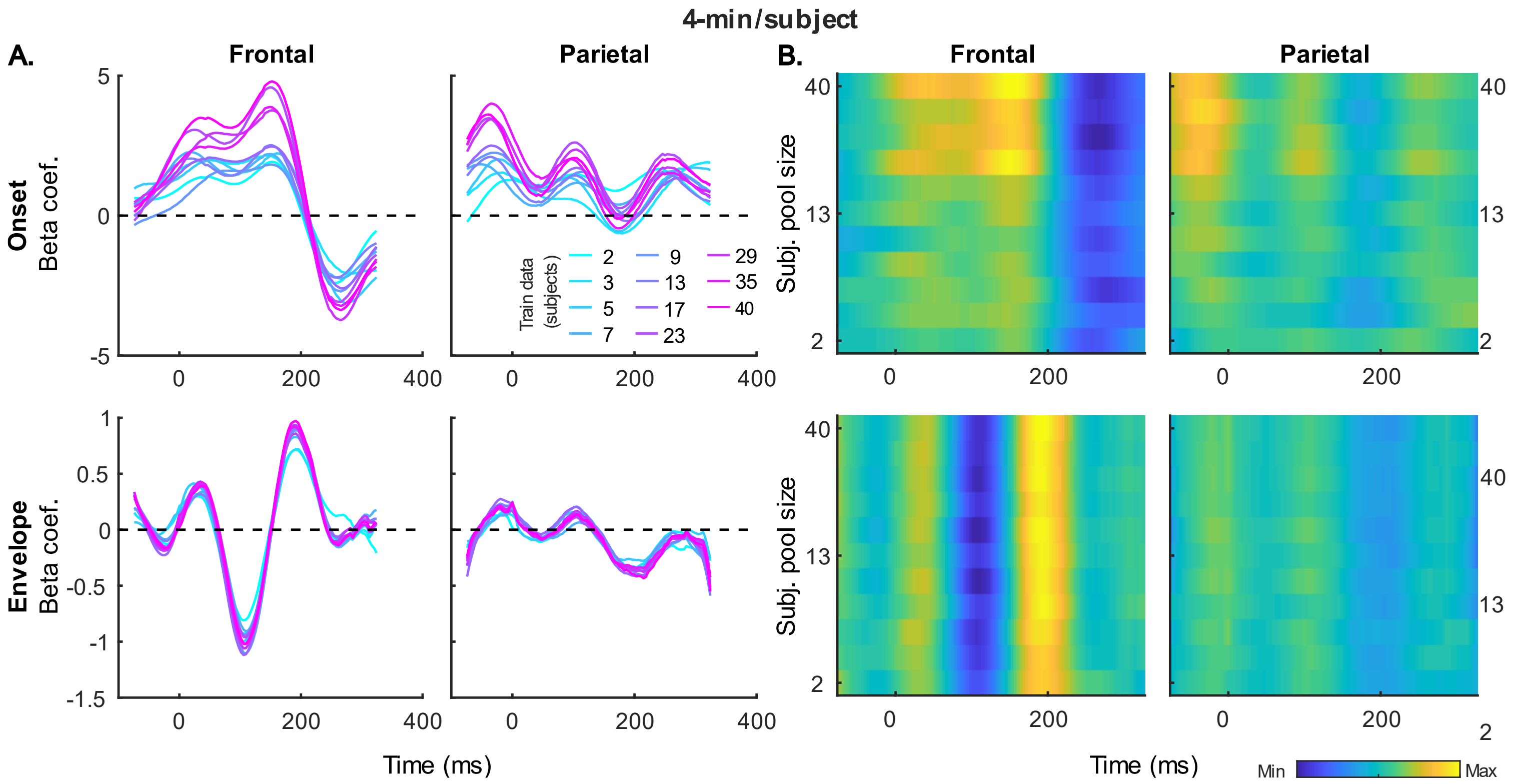


Supplementary Figure S6. Generic TRF time courses of the denser model trained using 4-min of data per participant. Same features and visualization conventions are used as in Fig S5.

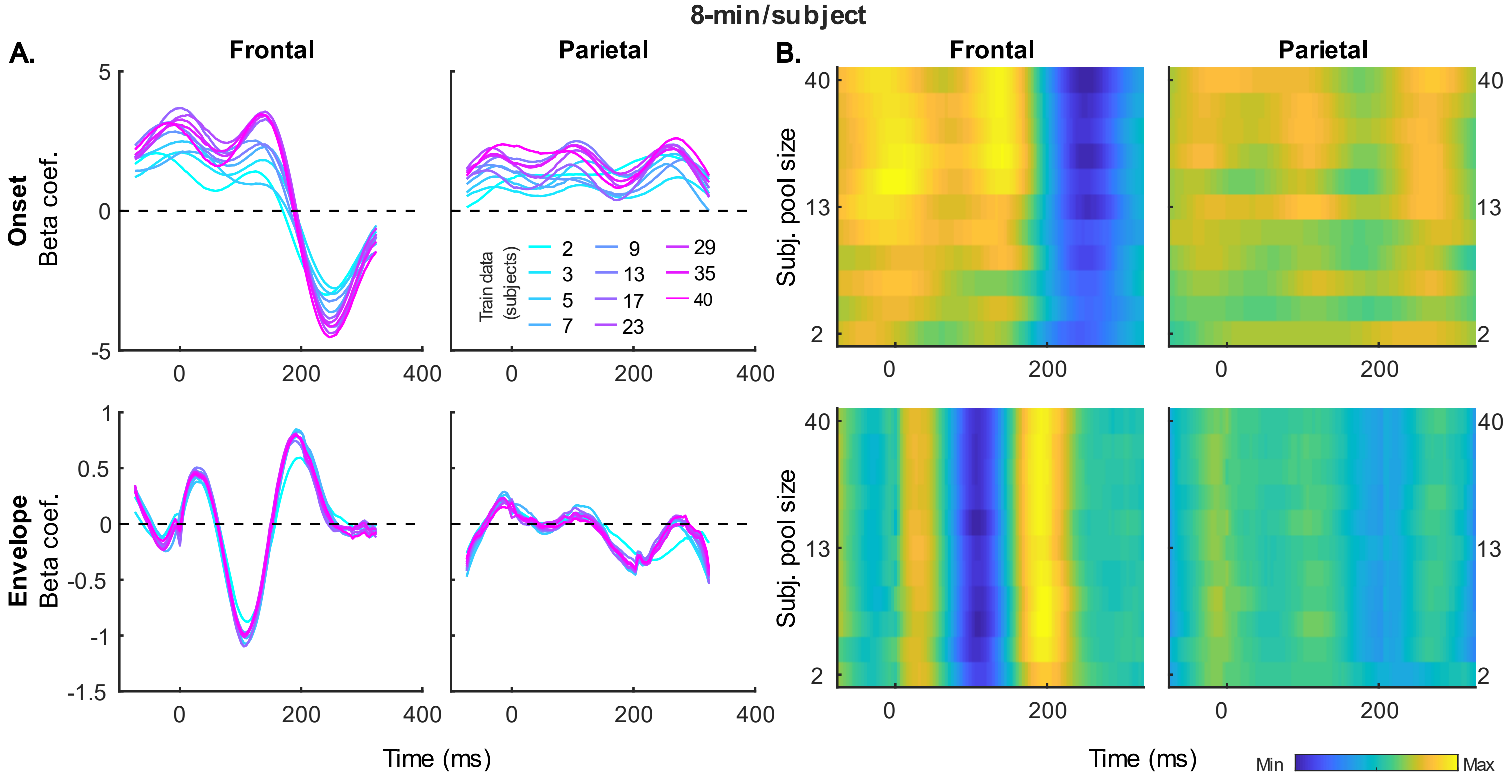


Supplementary Figure S7. Generic TRF time courses of the denser model trained using 8-min of data per participant. Same features and visualization conventions are used as in Fig S5.

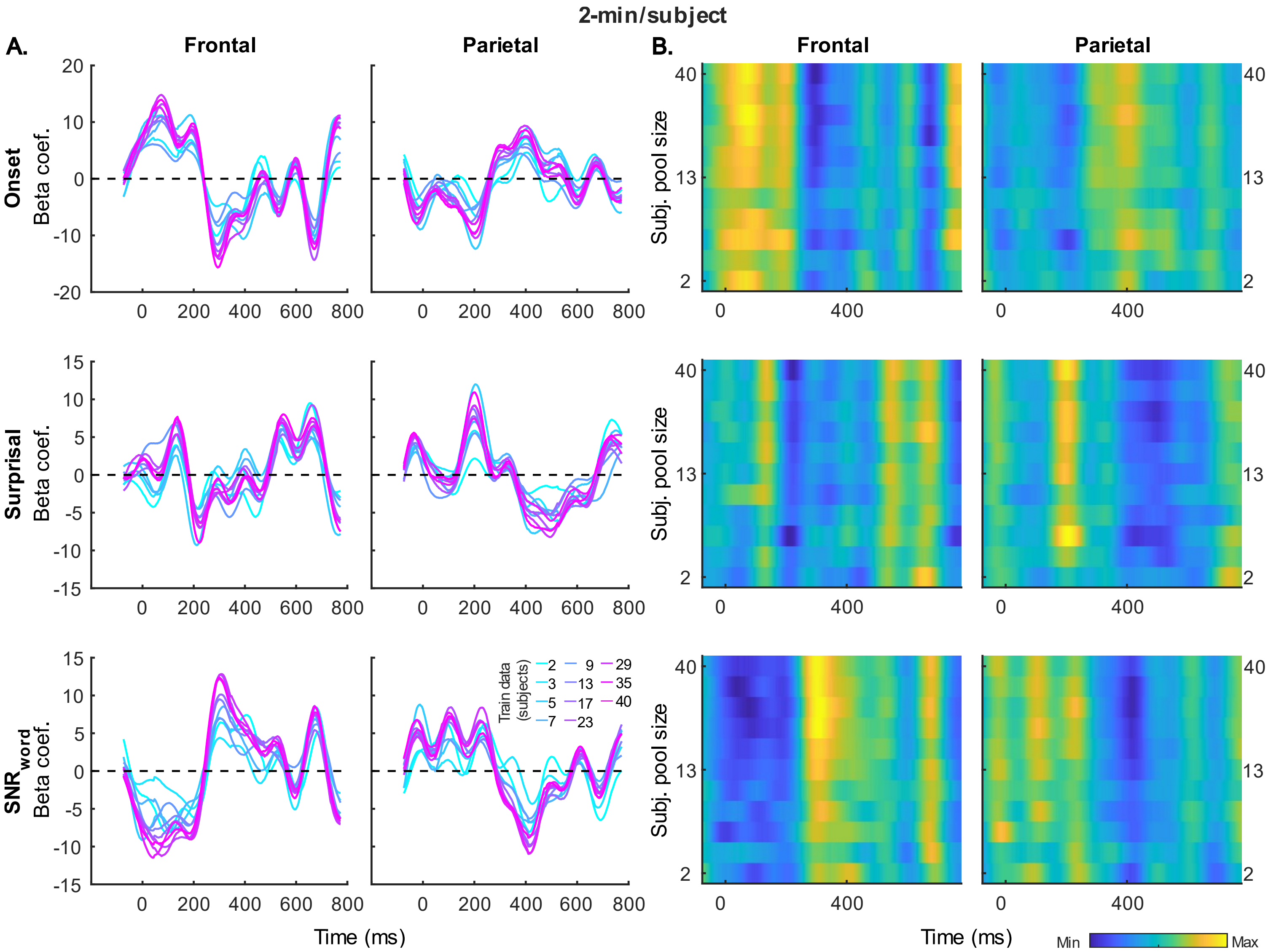


Supplementary Figure S8. Generic TRF time courses of the sparser model trained using 2-min of data per participant. Same visualization conventions are used as in Fig S5.

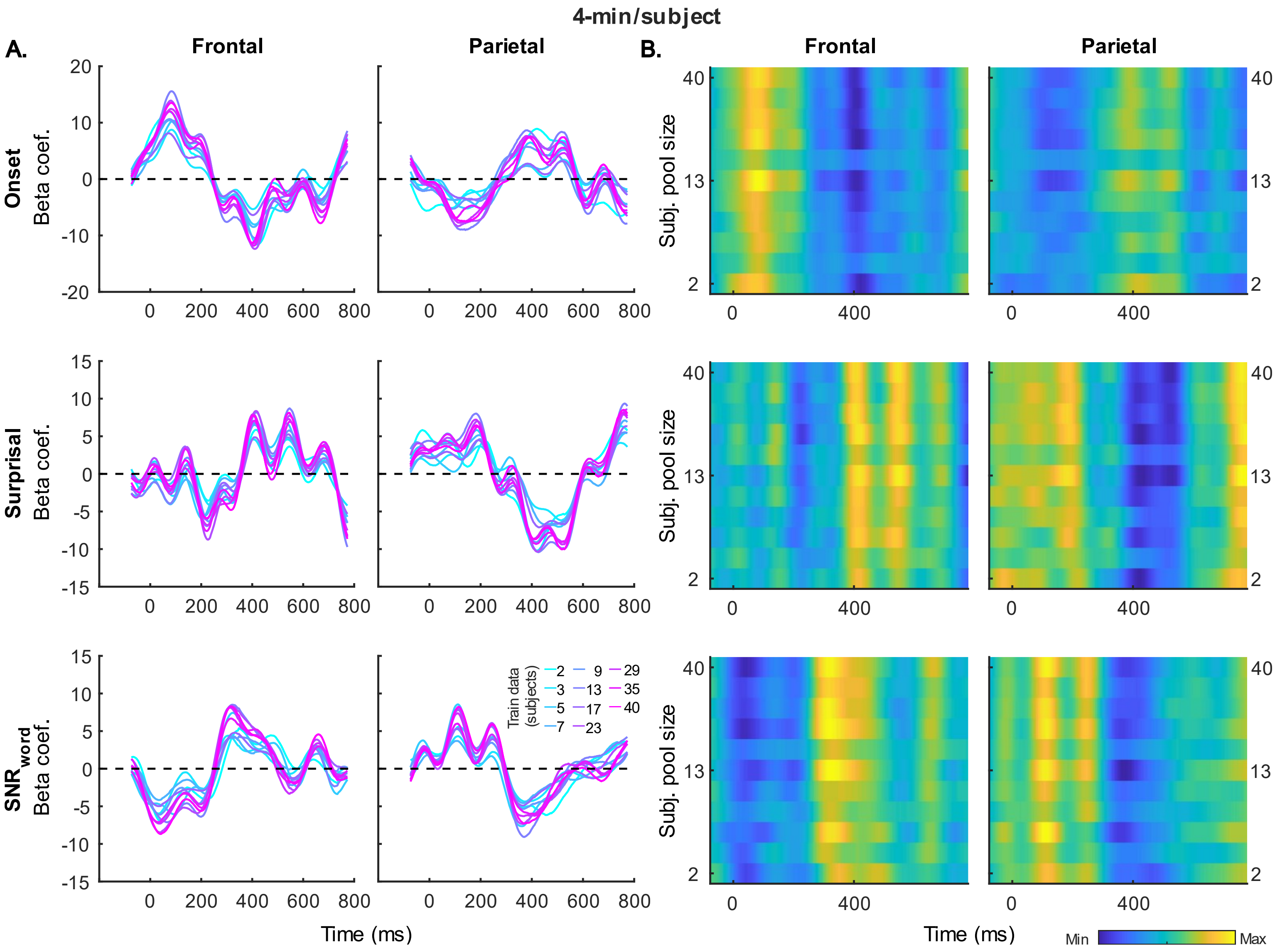


Supplementary Figure S9. Generic TRF time courses of the sparser model trained using 4-min of data per participant. Same visualization conventions are used as in Fig S5.

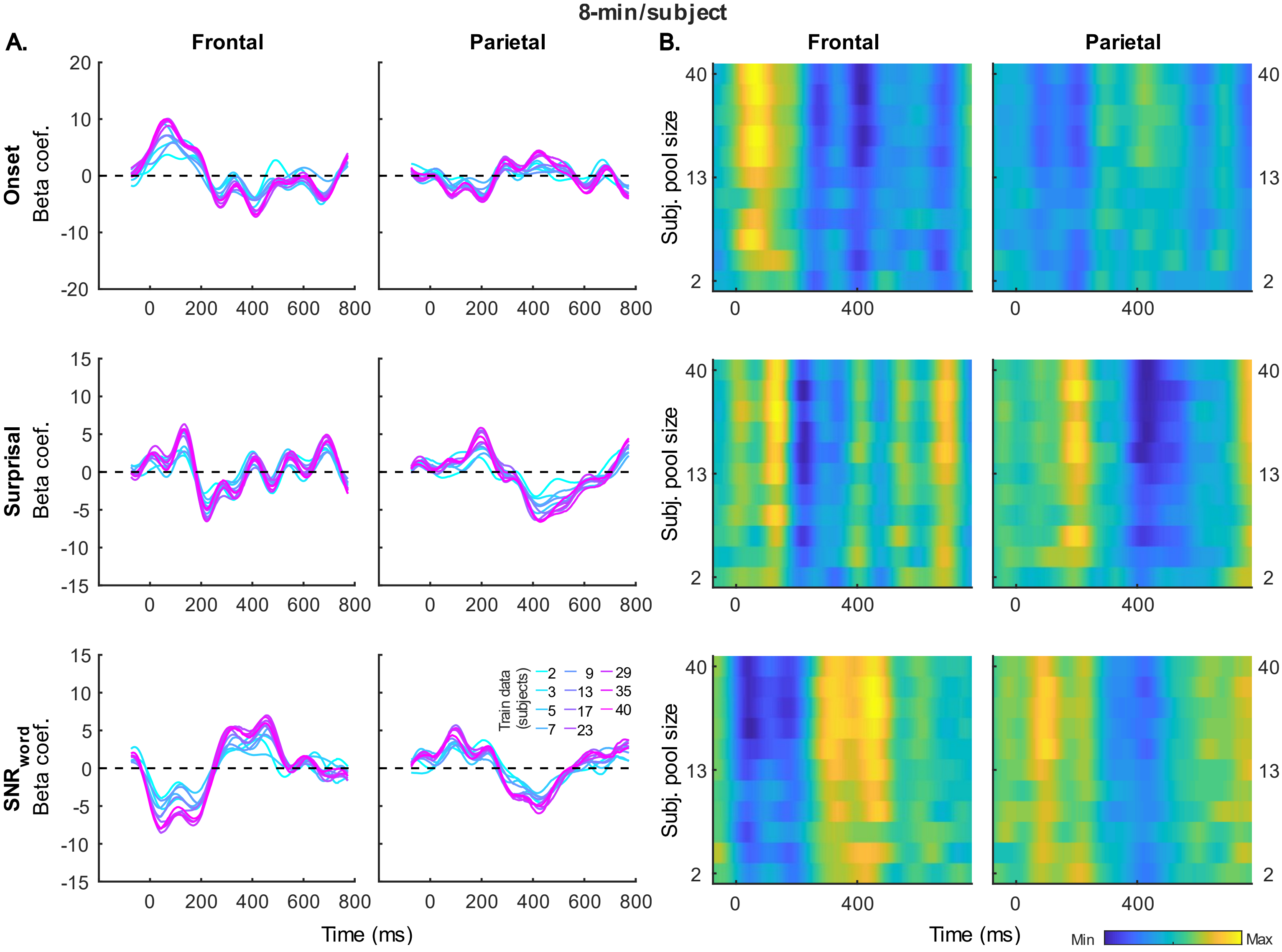


Supplementary Figure S10. Generic TRF time courses of the sparser model trained using 8-min of data per participant. Same visualization conventions are used as in Fig S5.

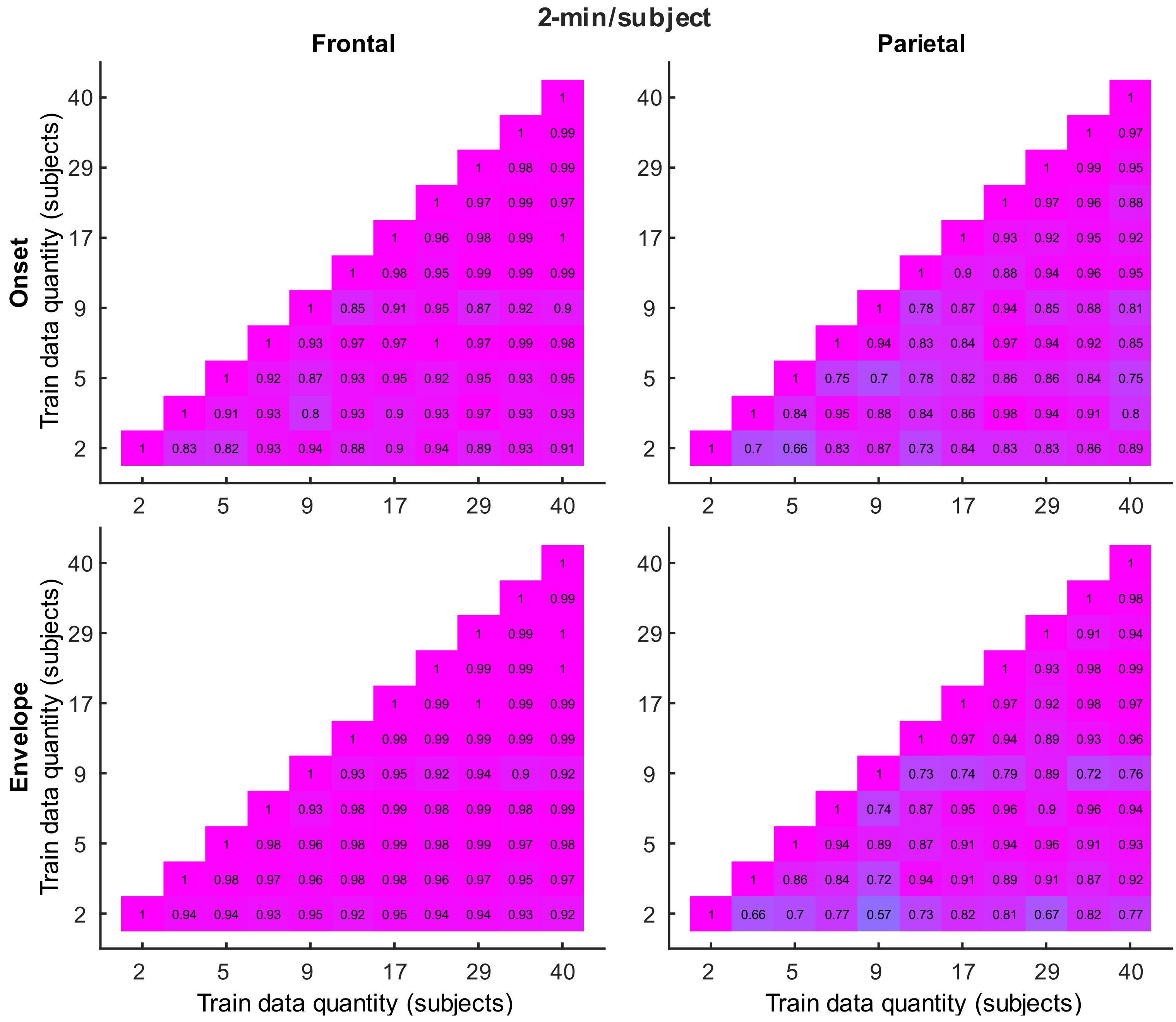


Supplementary Figure S11. Similarity of the denser, generic model TRFs derived using 2-min of data per participant for onset (top row) and envelope (bottom row) features in frontal (left column) and parietal (right column) ROIs. Each plot depicts pairwise Pearson's correlations between TRF time courses for all combinations of training data quantities (indicated along x and y axes of the plots). Brighter purple and blue colors reflect higher positive and negative correlation coefficients, respectively. Coefficient values are also indicated numerically at the center of each cell.

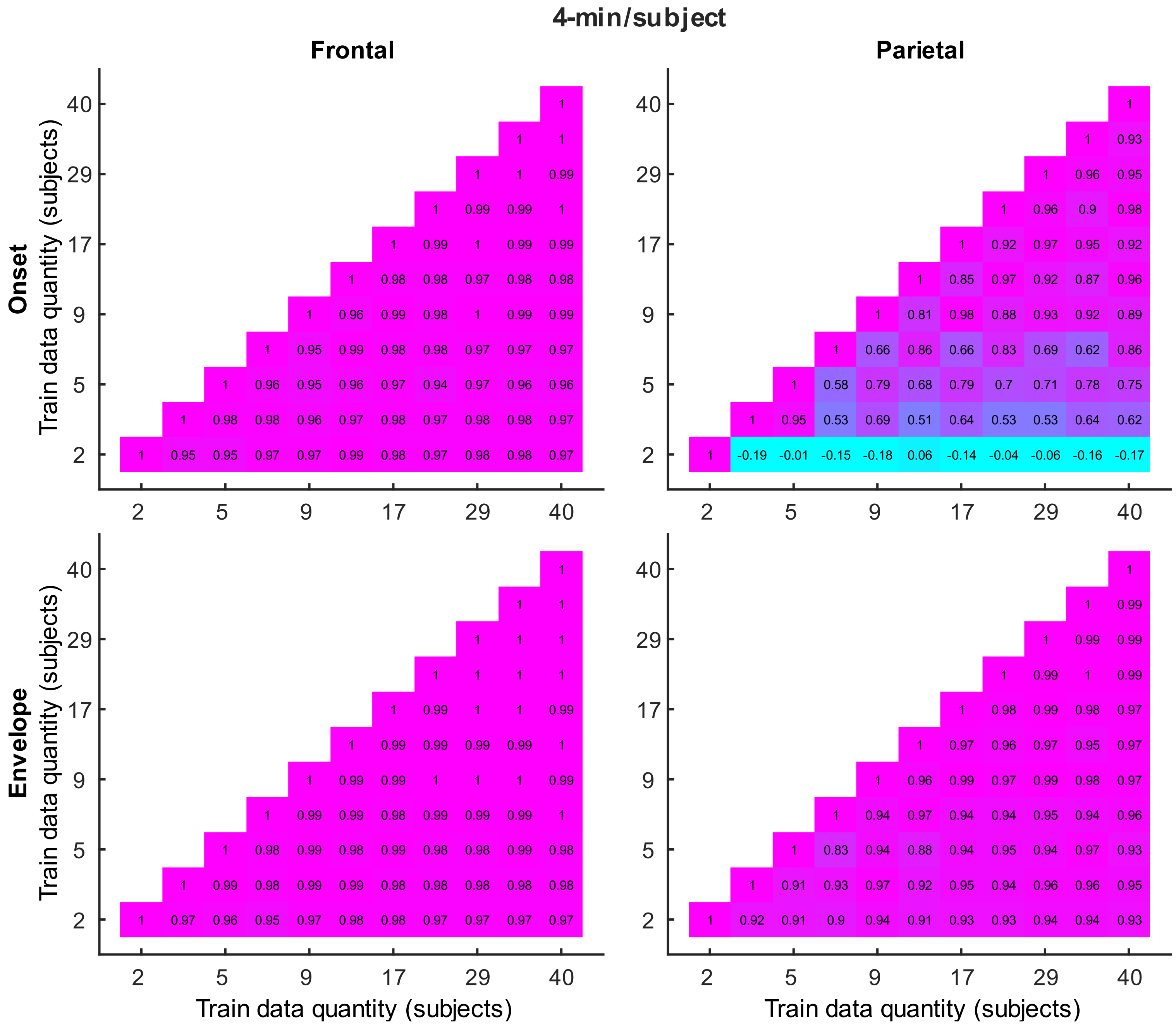


Supplementary Figure S12. Similarity of the denser, generic model TRFs derived using 4-min of data per participant. Same features and visualization conventions are used as in Fig S11.

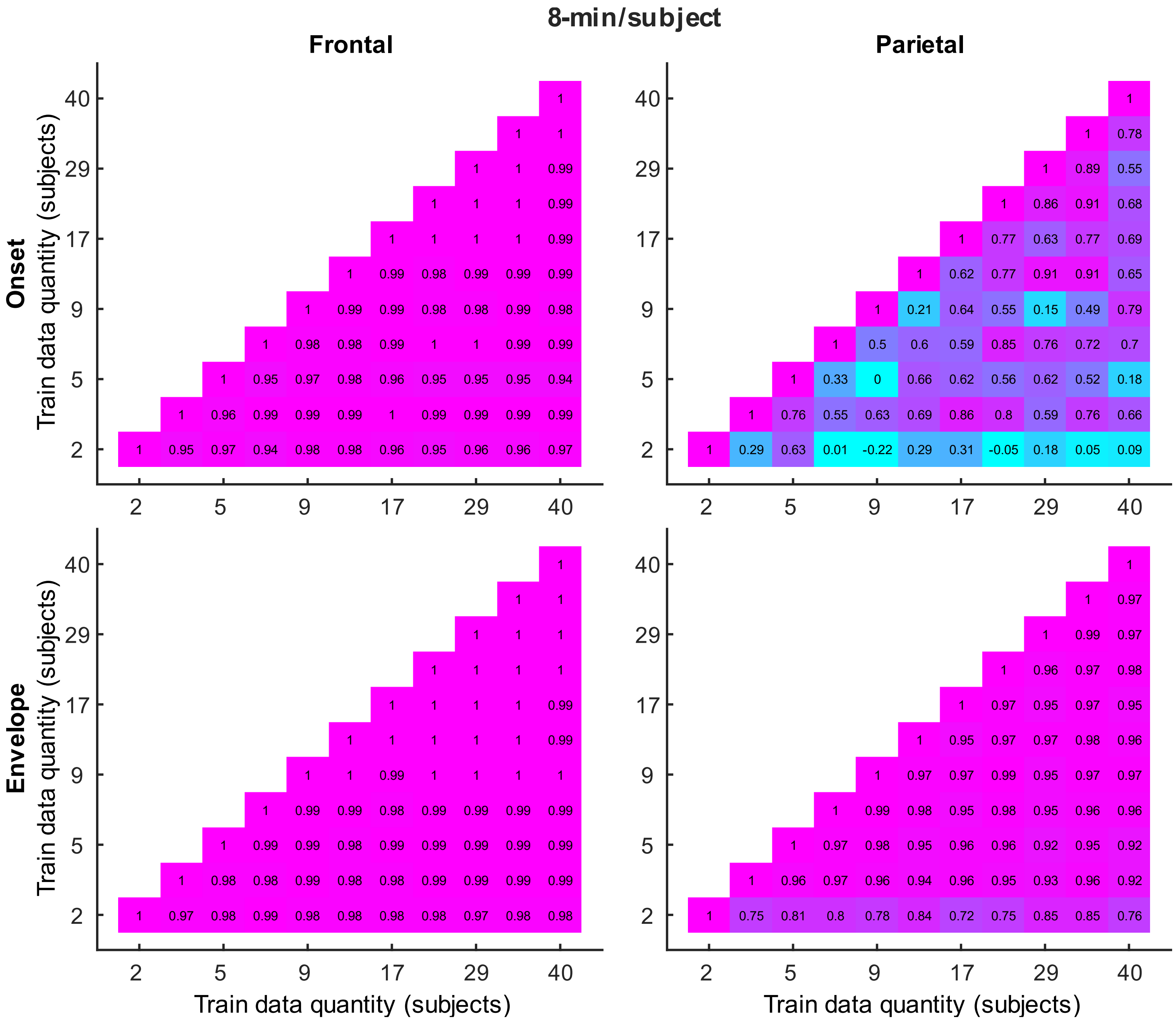


Supplementary Figure S13. Similarity of the denser, generic model TRFs derived using 8-min of data per participant. Same features and visualization conventions are used as in Fig S11.

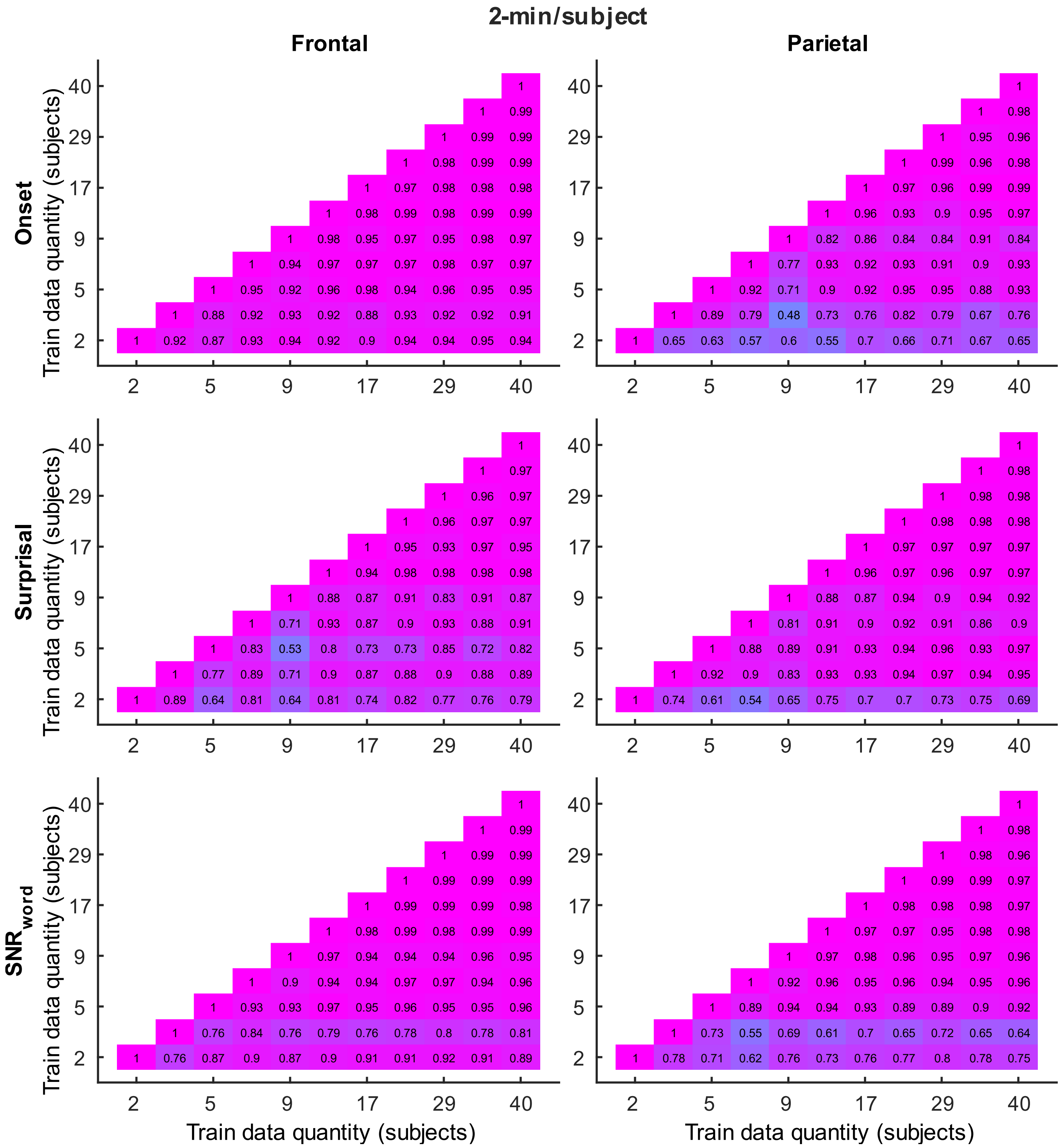


Supplementary Figure S14. Similarity of the sparser, generic model TRFs derived using 2-min of data per participant. Visualization conventions are used as in Fig S11.

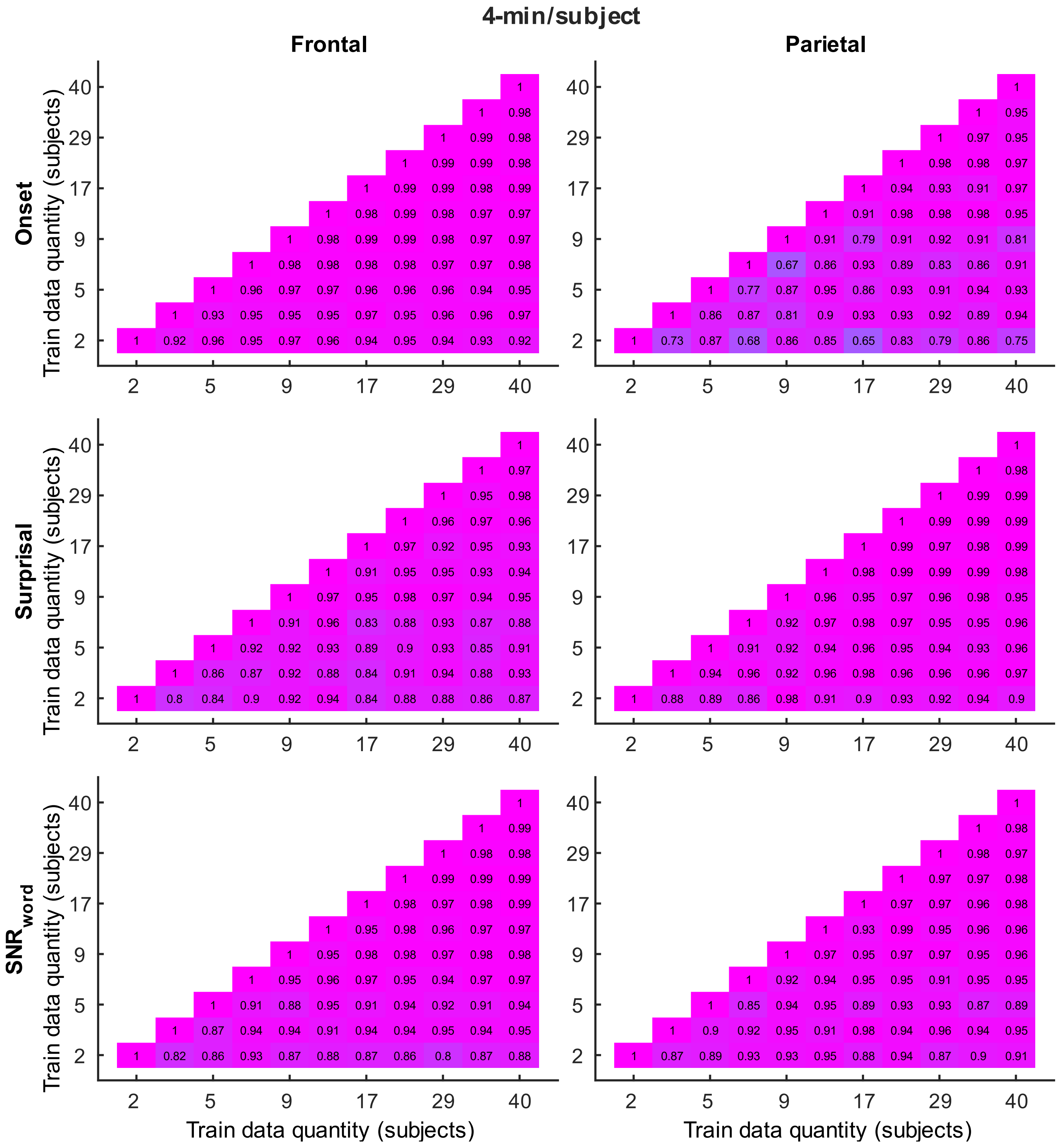


Supplementary Figure S15. Similarity of the sparser, generic model TRFs derived using 4-min of data per participant. Visualization conventions are used as in Fig S11.

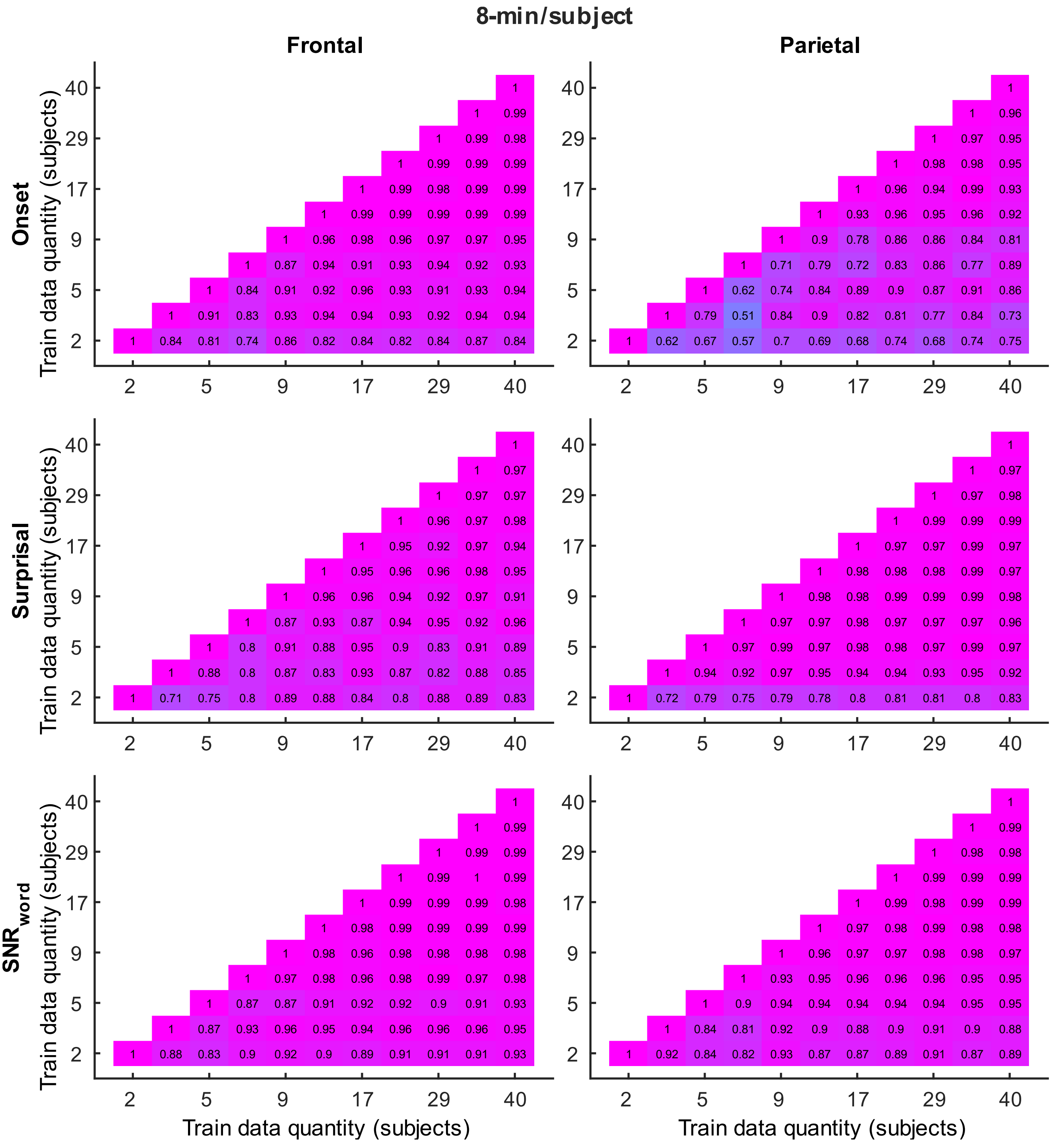


Supplementary Figure S16. Similarity of the sparser, generic model TRFs derived using 8-min of data per participant. Visualization conventions are used as in Fig S11.
